# Functional brain organization is stable within individuals across years

**DOI:** 10.64898/2026.06.10.731401

**Authors:** Hyejin J. Lee, Ally Dworetsky, Alexis Porter, Sihan Fei, Ashley N. Nielsen, Benjamin A. Seitzman, Babatunde Adeyemo, Patrick G. Bissett, Russell A. Poldrack, Jessica R. Cohen, Mark D’Esposito, Maital Neta, Nico U. F. Dosenbach, Steven E. Petersen, Evan M. Gordon, Deanna J. Greene, Caterina Gratton

## Abstract

Brain regions exhibit dynamic yet highly coordinated activity patterns that form large-scale functional networks measurable through resting-state correlations. While their association with fluctuating activity may intuitively suggest functional networks to be temporally transient and dependent on state, a growing body of literature suggests that they are person-specific and stable across days. If truly person-specific, then functional networks should preserve unique characteristics over extended periods.

To test this hypothesis, we collected longitudinal *precision* fMRI data (≥60 minutes per participant per time point) from 10 healthy young adults across 1-3 year intervals, as well as three adults over 8-13 years. We further replicated findings in the MyConnectome dataset and its 10-year follow-up. Functional network organization—when sufficient per-participant data were collected— remained largely stable within individuals over prolonged periods of up to 13 years, suggesting that individualized brain organization constitutes persistent features of personal identity that may be supported by homeostatic mechanisms.

**SIGNIFICANCE:** While many aspects of brain function are portrayed as dynamic and malleable, our study provides evidence for the long-term preservation of functional organization of the healthy young adult brain. Whole-brain functional organization exhibited unique individual characteristics that were preserved for years, even over a decade. The stability of such individualized neural architecture suggests that the brain encodes features of personhood that may be invariant across contexts and temporal fluctuations. This observation reframes our understanding of brain resilience and plasticity by offering new insights into the extent that homeostatic mechanisms of brain organization can withstand stressors and environmental changes through time to maintain a constant functional organization. These insights pave the way for a more comprehensive understanding of the neural mechanisms behind this stability, with the possibility of pinpointing critical markers of healthy brain function and its disruptions.

## INTRODUCTION

Many aspects of brain function are portrayed as fluctuating, dynamic, and malleable depending on internal states and environmental demands. Yet, people are typically stable in their cognitive abilities and behavioral profiles across adulthood (Costa, 1986; Graham et al., 2020; Schaie et al., 2004), contributing to a person’s sense of stable identity. A fundamental question is what aspects of brain function persist similarly unchanged, supporting stable, trait-like aspects of personhood.

Recent advances in fMRI have made it possible to examine functional organization in the human brain in vivo in a replicable manner (Power et al., 2011; Thomas Yeo et al., 2011). A characteristic aspect of this organization is the presence of distinct, large-scale systems, or intrinsic ‘functional networks’, of brain areas that show correlated activity, even while a person is at rest (Biswal et al., 1995; Damoiseaux et al., 2006; Fox & Raichle, 2007; Smith et al., 2009). Although there are general principles shared across all brains, these investigations have highlighted that no two brains mirror each other’s functional architecture (Finn et al., 2015; Miranda-Dominguez et al., 2014): significant inter-individual variability exists, giving rise to unique, idiosyncratic features in functional networks (Dworetsky et al., 2024; Gordon et al., 2017; Seitzman et al., 2019). These individual characteristics can predict variation in behavior and cognition among individuals (Bijsterbosch et al., 2018; Finn et al., 2015; Kong et al., 2019, 2021; Lee et al., 2025; Pierce et al., 2024), suggesting functional networks may encode person-specific traits.

A fundamental question raised by these results is whether functional networks remain stable across adulthood, even over extended periods marked by stressors and environmental changes. This pattern would support the idea that homeostatic mechanisms maintain functional network integrity (Laumann & Snyder, 2021) and, in turn, core phenotypic characteristics. One theoretical account proposes that inter-regional correlations in brain activity reflect cumulative history of brain activity accrued over an individual’s lifetime (Petersen & Sporns, 2015). Within this framework, homeostatic mechanisms may act to preserve functional network integrity despite ongoing context-driven neural fluctuations (Laumann & Snyder, 2021). This perspective argues that individual-specific functional organization should be stable across different mental states and over prolonged periods within individuals, thereby preserving individual identity. Consistent with this, prior work has demonstrated that functional network organization remains largely stable within individuals over days and weeks (Gordon et al., 2017; Laumann et al., 2015; Seitzman et al., 2019), including in the face of changing cognitive demands (Du et al., 2025a; Gratton et al., 2018).

However, it is not known how extensively the statistical history of functional brain activity might be altered across extended timescales, such as periods over a decade. Over these time windows, adults accrue changes in life experiences, stressors, and environmental input—yet their sense of identity remains intact. Extended longitudinal data may help to disambiguate between competing models for functional brain organization: whether it is shaped primarily by recent experiences or preserved to reflect trait-like features of personhood through homeostatic mechanisms. Although some research has suggested that at least a subset of large-scale brain networks can show some degree of individual identifiability over 1-3 years (Badhwar et al., 2020; Horien et al., 2019; Jalbrzikowski et al., 2020), others have highlighted dynamic variation in brain networks even within a single scan session (Albert et al., 2009; Allen et al., 2014; Dhond et al., 2008; Lurie et al., 2020; Sami & Miall, 2013).

Resolving this uncertainty requires high quality, longitudinal measures of functional brain organization at the individual level. Prior longitudinal studies often faced two major limitations. One is limited individual data—often less than 10 minutes per time point—where sampling variability critically constrains the reliability of fMRI measures (Elliott et al., 2019, 2020; Gordon et al., 2017; Laumann et al., 2015; Noble et al., 2017). The other is limited time spans of 1-3 years, not well matched to the expected windows of life experience change in adulthood. To address these limitations, our study employed high-quality, densely sampled data for each participant (≥60 minutes collected per time point) across a broad time range (1-13 years) to map individual network topography with high precision. We replicated our findings with the MyConnectome dataset, an extensively sampled, single-participant dataset originally released in 2014 (Laumann et al., 2015; Poldrack et al., 2015), and recently expanded with the addition of longitudinal data collected a decade later.

## RESULTS

We tracked precise measures of brain organization in the same individuals across an extended time interval: two participants were tracked across an interval of 13 years and one across an interval of 8 years (**Prolonged Longitudinal Cohort**; see **Fig. 1** for a summary of data collection). This cohort underwent significant life transitions, including childbirth, parenting, career changes, relocation across the country, loss of a parent, and pandemic lockdowns. We additionally acquired data from 10 individuals scanned across shorter intervals of 1-3 years with an identical scanning protocol (**Standard Longitudinal Cohort**). Our initial analyses focused on the Prolonged Cohort to evaluate the stability of network measures over extended periods.

**Figure. 1.**
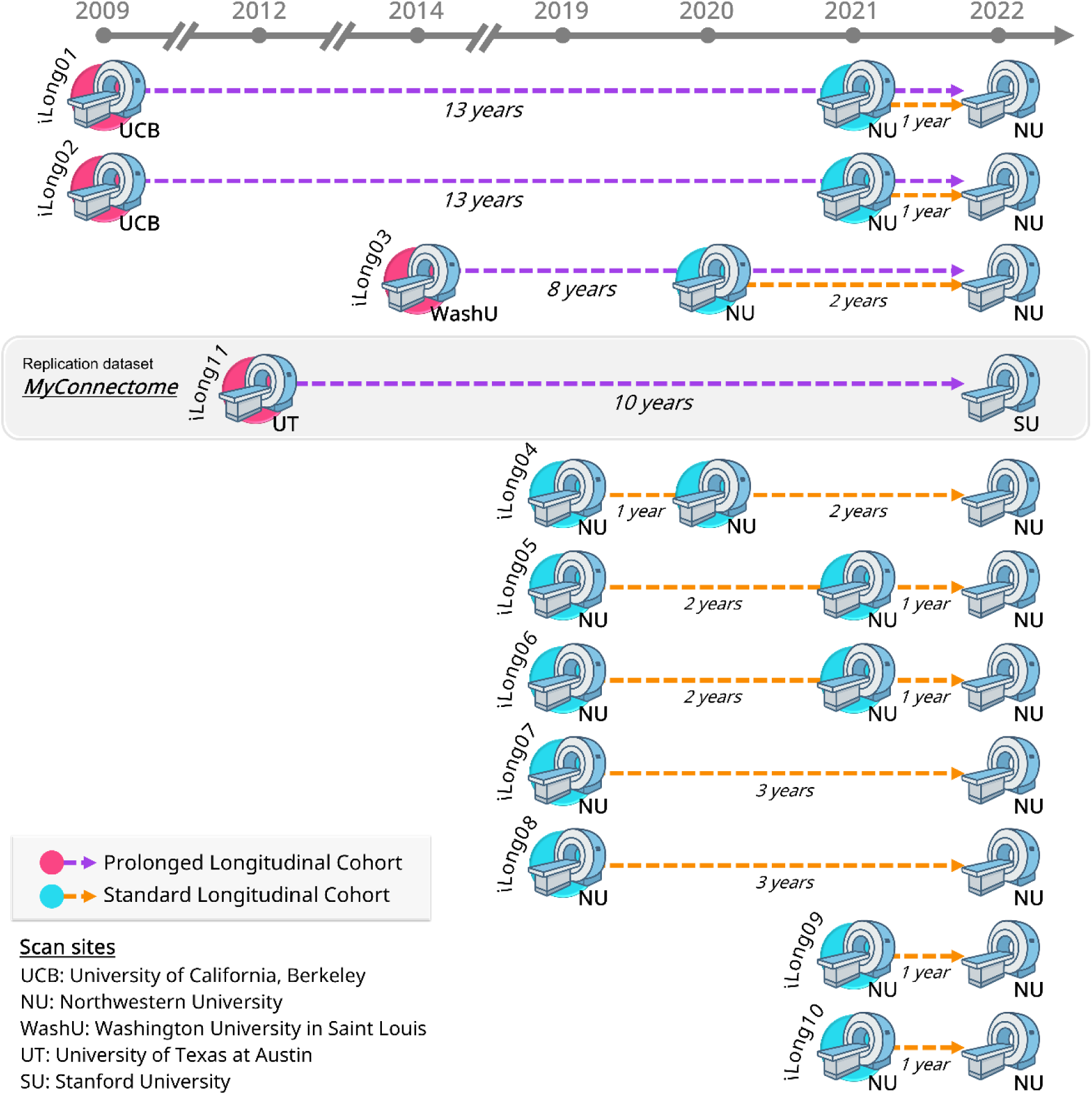
Data summary. We collected data from 10 individuals (iLong01-10; mean age of 26 years at the time of first scanning) and grouped into two cohorts, based on scan intervals and condition matching (prolonged and unmatched versus standard and matched). The Prolonged Cohort had extended intervals of 13 (iLong01 and iLong02) or 8 years (iLong03), with initial scans conducted at University of California, Berkeley (iLong01 and iLong02) and Washington University in Saint Louis (iLong03). The MyConnectome dataset (45 years old) and its follow-up were included as replication data (iLong11) to the Prolonged Cohort; here time point 1 was collected at the University of Texas at Austin, and time point 2 at Stanford University, 10 years later. The Standard Cohort had shorter intervals between scans of 1-3 years, with all scans performed at Northwestern University using a consistent protocol. More details on data acquisition can be found in the *Methods* section.

### 1.1 Functional organization exhibits within-individual stability over a decade-long period

In the three Prolonged participants, the organization of functional brain networks, individually mapped at each time point, was largely stable over extended periods (**Fig. 2a**). Consistent with prior research (Gordon et al., 2017), brain network topography was unique in each participant (comparing each participant’s first time point with the group average in **Fig. 2a**). Notably, within each participant, functional network topography was largely stable over 8-13 years. Most vertices (locations on the cortical surface) were assigned to the same network at both time points (79% for iLong01 and iLong02, and 83% for iLong03), significantly exceeding the null distribution derived from the between-participant comparisons (*M* = 64%, *SD* = 1.65; permutation *p* < 0.001; analyses conducted after excluding low-SNR regions). These results were replicated in the MyConnectome dataset and accompanying longitudinal data collected 10 years later (75% consistency across time points; **Fig. 2a**, bottom row). Notably, in all four participants, even small, idiosyncratic network regions (Dworetsky et al., 2024)—unique to each participant and not observed in the group average—remained consistent over time. We highlight examples within the lateral prefrontal area (**Fig. 2a**, right zoom-in) where some of the largest individual variability in network topography has been observed (Ladwig et al., 2026).

**Figure 2.**
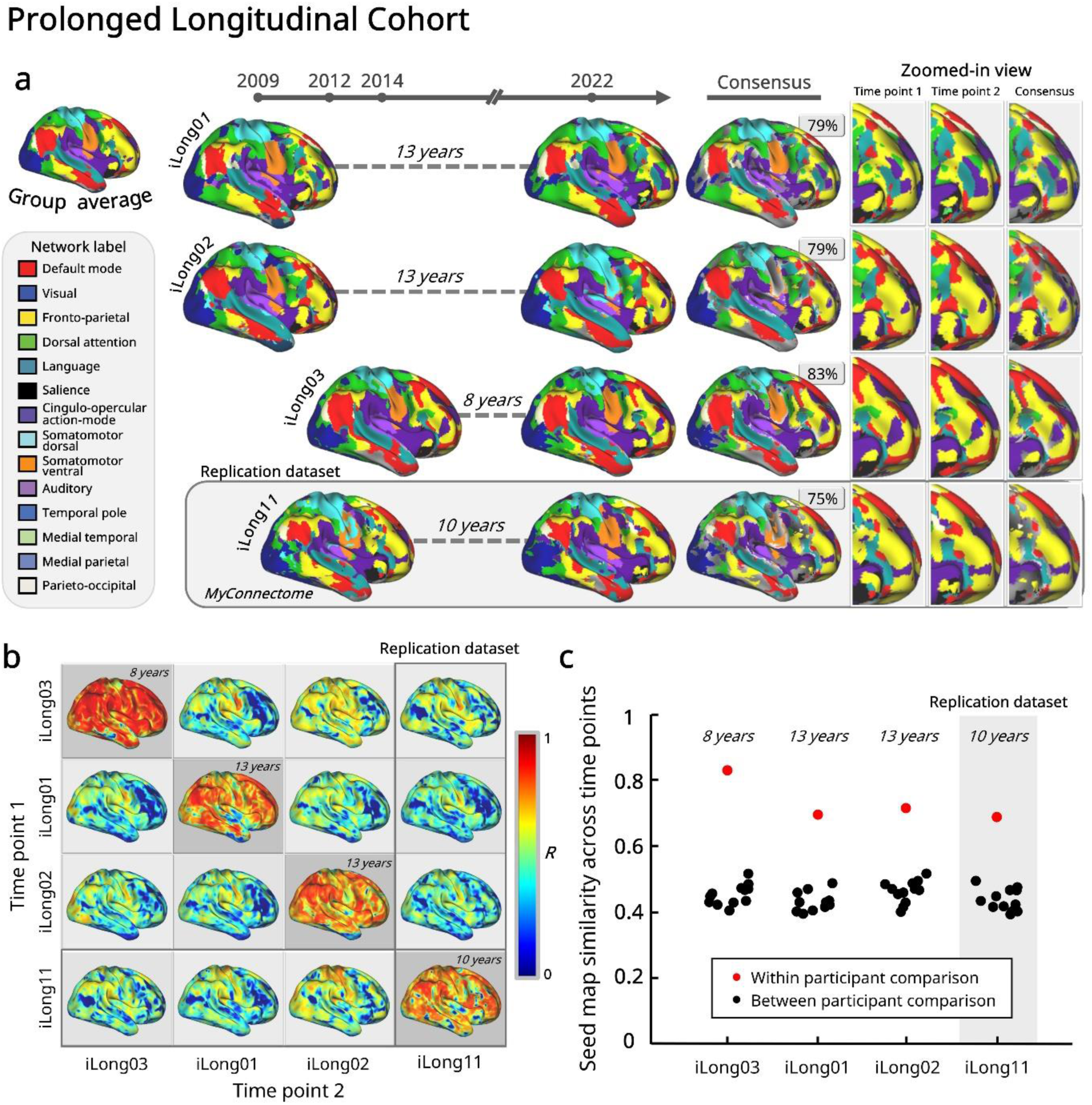
Individual-specific functional brain organization is stable over prolonged periods. **a) Network maps show high temporal consensus.** Large-scale functional networks at each time point are shown with colors indicating 14 distinct networks. Results are shown separately for each of the Prolonged Longitudinal Cohort (rows) in chronological order. The final consensus map shows color-coded vertices assigned to the same network across both time points (inconsistent vertices are shown in gray). Results show high consensus in all three participants, as well as in the replication dataset. A zoom-in highlights the consistency of individual specific regions in the lateral prefrontal cortex. **b) Continuous seed maps replicate high temporal stability.** Each vertex is colored by the similarity (Pearson’s *r*) of functional connectivity seed maps for each vertex between prolonged time points. High consistency is seen within the same participant over time (diagonal spaces), whereas the similarity between participants was notably lower. This pattern was consistent in the replication dataset (last row/column). **c) Within-participant stability significantly exceeds between-participant variability.** The average of the whole-brain maps presented in panel b with low-SNR regions censored. Red dots represent the average similarity (*r*) across time points within participants, which were significantly higher than the between-participant correlations shown in black.

To provide a more detailed picture, whole-brain functional connectivity profiles for each vertex were compared across time points (**Fig. 2b**). Again, the results showed robust within-participant stability; the average spatial correlation across the whole cortex, calculated across the three Prolonged participants, was *r* = 0.75 (*SD* = 0.07), which was significantly higher than the functional connectivity similarity between participants (*r* = 0.45, *SD* = 0.03) with permutation testing, *p* < 0.001 (**Fig. 2c**). This pattern was consistently observed in the replication dataset (across time points, *r* = 0.69). Regions with relatively lower stability primarily fell into areas of lower signal quality (e.g., inferior temporal and orbitofrontal; (Ojemann et al., 1997a)) or regions known to be more susceptible to arousal-related effects (visual and motor; (Laumann et al., 2015; Tagliazucchi & Laufs, 2014)). Note that this level of stability was achieved despite the fact that scanners and scan sequence parameters were not matched across time points. As such, consistency across these datasets should be considered a lower bound of the similarity that functional brain organization can achieve across these timescales.

### 1.2 Controlled scanner conditions show comparable stability across weekly and yearly timescales

We subsequently conducted systematic investigation of longitudinal changes in 10 participants (**Standard Longitudinal Cohort**) with data collected on the same scanner with the same scan parameters over 1-3 years (see *Methods*). Critically, this dataset features multiple sessions at each of 2-3 time points, allowing for a comparison between the amount of day-to-day variability between sessions relative to variability across years.

Results showed remarkable stability in functional network organization. Across 10 participants, 87% of vertices (*SD* = 3.91) were assigned to the same network across all longitudinal time points (**Fig. 3a**). This degree of network consistency was largely stable across temporal scales (**Fig. 3b**, based on matched data quantities), with similarity both for sessions separated by weeks (*M* = 82%, *SD* = 2.93) and years (*M* = 81%, *SD* = 3.37) significantly exceeding between-participant similarity (black; *M* = 61%, *SD* = 1.65), *p* = 0.002 (group-level permutation testing via label shuffling).

**Figure 3.**
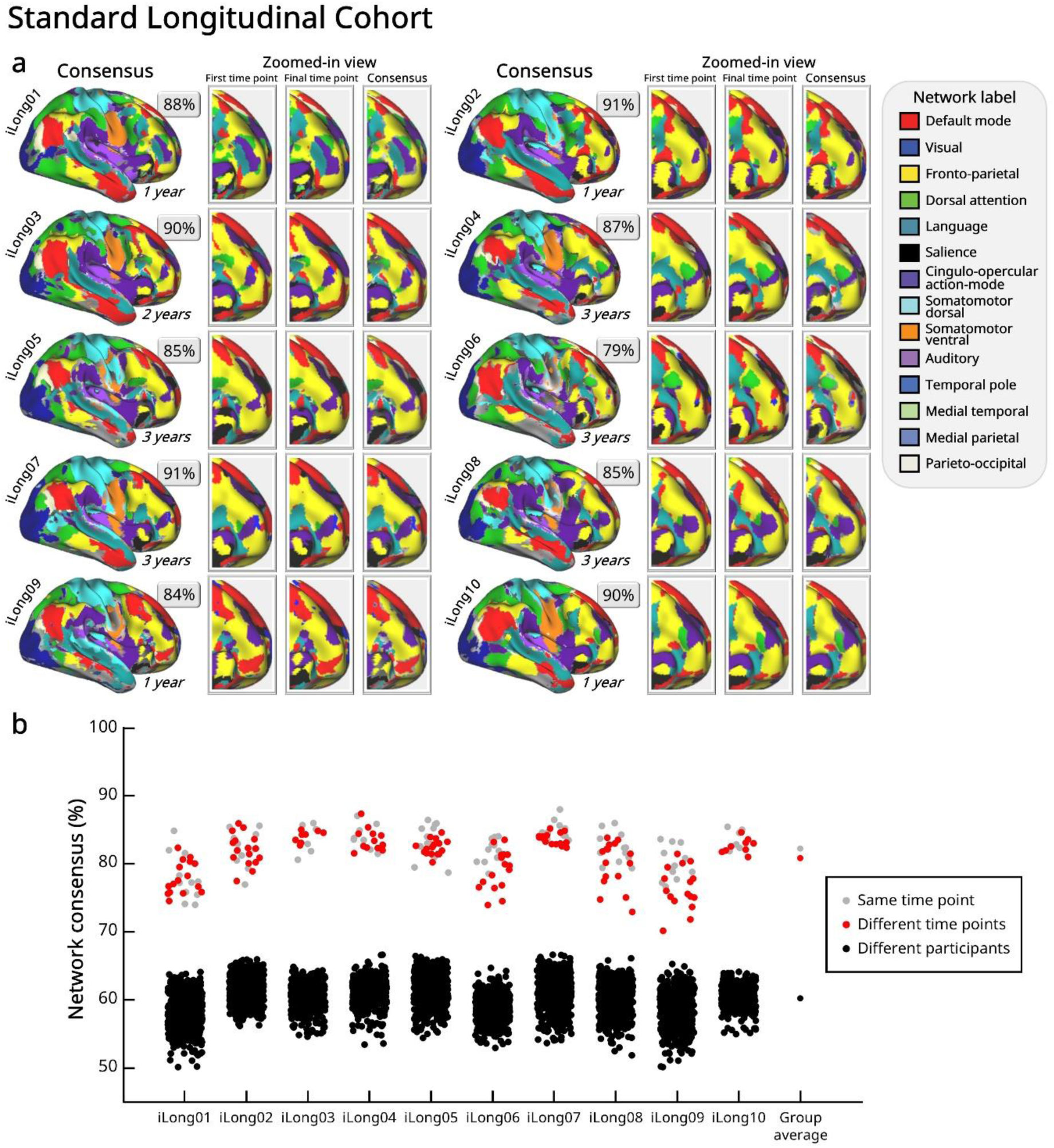
Controlled measurements reveal strong stability of functional network organization. **a) High network consensus with preserved idiosyncratic features.** We mapped functional networks at the individual level and colored the vertices when the same network was assigned across all two or three time points in the Standard Longitudinal Cohort (as in Fig. 2a; time delay shown under each brain). With controlled measurements, most vertices were consistent over time. The zoomed-in view shows idiosyncratic locations in the lateral prefrontal cortex, which vary across participants but appear consistently within participants across time points. See **Supp. Fig. 1** for the full timeline and brain maps. **b) Within-participant similarity remains high across same and different time points relative to between-participant comparisons.** As expected, while similarity was greatest across weeks (same time point; gray dots), similarity across years (different time points; red dots) also remained high and was significantly greater than between-participant stability (black dots). The data quantity was matched (30 minutes per session), and low-SNR vertices were censored.

As before, we also examined the similarity in continuous maps of vertex-wise functional connectivity pattern (**Fig. 4**), without enforcing a hard network parcellation (as in **Fig. 3a**). Similarity across time points was high throughout the cortex (**Fig. 4a**), with a participant group mean of *r* = 0.91 (*SD* = 0.03, excluding low-SNR regions). Comparisons with matched data across participants (86 minutes per time point; **Fig. 4a** matrix) confirmed that the within-participant similarity (*r* = 0.88, *SD* = 0.02) was almost double the between-participant similarity (*r* = 0.48, *SD* = 0.04; permutation test, *p* < 0.001). Likewise, comparisons across sessions (30 minutes per session, separated across weeks or years; **Fig. 4b**) confirmed that within-participant correlations (*M* = 0.77, *SD* = 0.05) remained consistently high, whereas between-participant correlations (*M* = 0.4, *SD* = 0.04) were significantly lower, *p* < 0.001, recapitulating the pattern observed across yearly time points.

**Figure 4.**
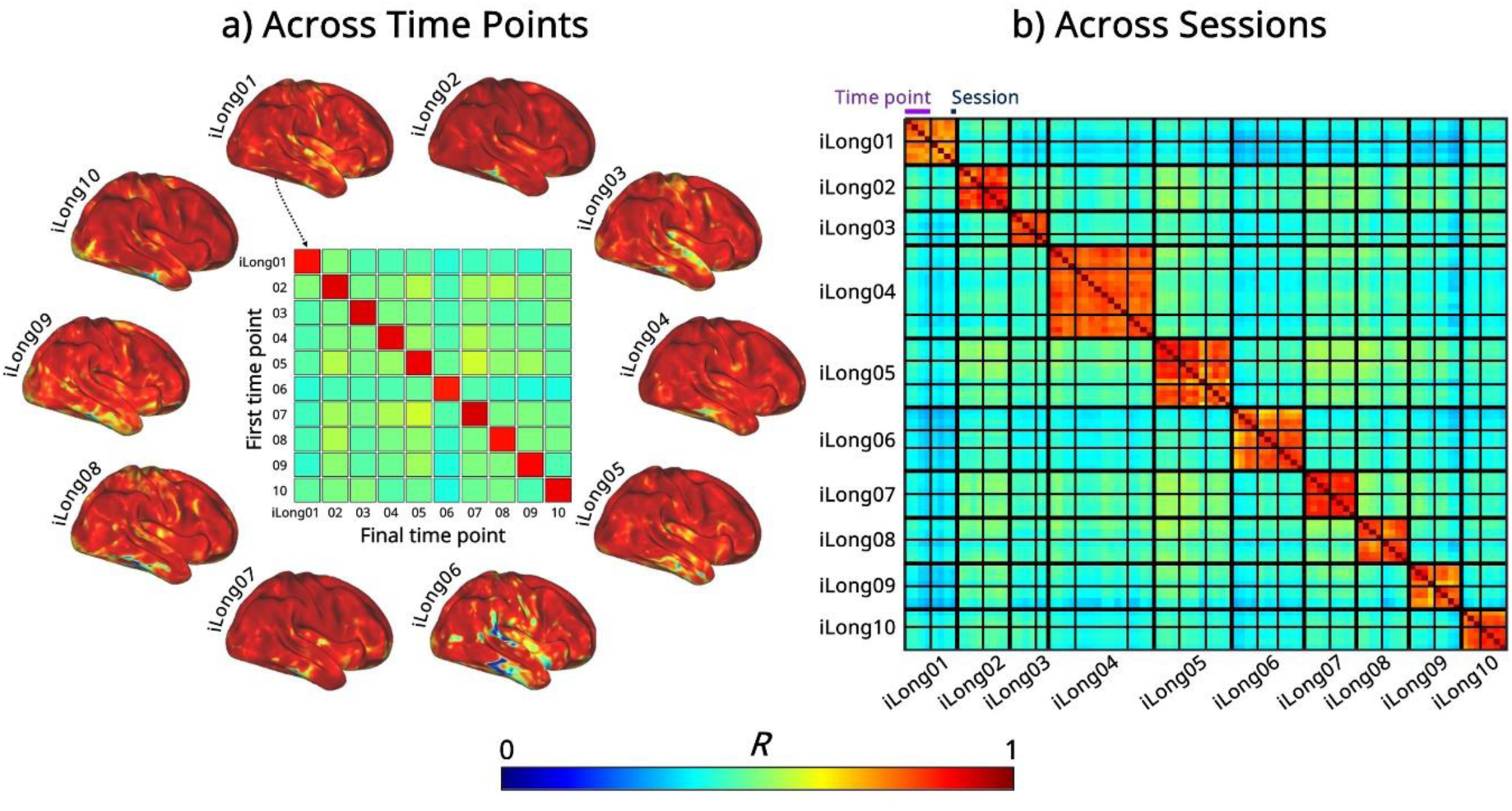
Robust within-participant stability in functional connectivity for both yearly and weekly data. **a) Stability across yearly time points.** The spatial distribution of functional connectivity similarity across the brain in the 10 Standard Cohort participants. For each participant, functional connectivity seed maps between the longest intervals (1-3 years) were compared for each vertex. The results showed high spatial correlations, *r* > 0.8 for most throughout the brain. The central matrix shows within-participant comparisons along the diagonal, which were significantly higher than the off-diagonal between-participant comparisons. The matrix values represent average spatial correlation across the cortical surface after censoring low-SNR regions. **b) Stability across sessions.** The session-level correlations were highly stable both within (weekly) and across (yearly) time points but differed substantially across participants; note that the correlations were comparable within participants (denoted by the thickest black lines) across different time points (next thickest black lines), as reflected by the warm color gradients along the diagonal, consistent with the matrix in panel a.

Consistent results were seen with regional analyses done in the cortical volume instead of on the cortical surface to ensure that findings are robust irrespective of analysis framework (**Supp. Fig. 2**). These results recapitulate that in young adults, functional correlation patterns are relatively stable across years, with yearly variability only subtly exceeding variability explained by measurement noise and daily fluctuation.

To more systematically assess the influence of elapsed time on functional connectivity similarity, we formally modeled inter-session similarity as a function of elapsed time between sessions (days), signal quality (BOLD tSNR), data quantity (frames remaining after motion censoring), and participant baseline, using a linear fixed effects model for each region (Seitzman300 atlas). Based on the observations above, we hypothesized that elapsed time would account for only a small proportion of variance. Global variance partitioning (**Supp. Fig. 3a**) revealed that the modeled factors accounted for approximately 30% of the variance in functional connectivity similarity across sessions. This relatively low proportion of explained variance is attributable to high baseline functional connectivity similarity in most regions, as indicated by the high mean intercept of 0.9 (**Supp. Fig. 3b**). Within the explainable portion of the variance, participant baseline was the dominant contributor (24.75%), indicating inter-individual differences in baseline similarity, whereas elapsed time (2.58%), BOLD signal quality (0.54%) and data quantity (1.55%) accounted for only small fractions (note however that data quantity was large and relatively similar across these datasets). Thus, differences in elapsed time (weeks to years) had only a small (2.6%) effect on similarity among functional network maps.

### 1.3 Precise estimates of longitudinal stability require sufficient within-participant data

Sampling variability arises from limited amounts of fMRI data, leading to fluctuations that do not reflect true neural changes. To demonstrate the significance of reducing sampling variability in longitudinal assessment, we compared functional connectivity derived from high- and low-sampling regimes. We compared functional network similarity derived from precision fMRI data (30 minutes per session) relative to more conventional non-precision data collection protocols where sampling variability is high (matched 10 minutes of low-motion data; **Fig. 5**). Our findings of robust stability across sessions were stronger with high precision sampling (within-participant similarity across sessions of *r* = 0.77) than with smaller amounts of data (within-participant similarity of *r* = 0.62), as sampling variability significantly weakened the similarity across sessions, *p* < 0.001 (permutation testing). This highlights the added consistency in individual brain network measures that can be achieved with extended multi-session scanning, relative to more standard protocols.

**Figure 5.**
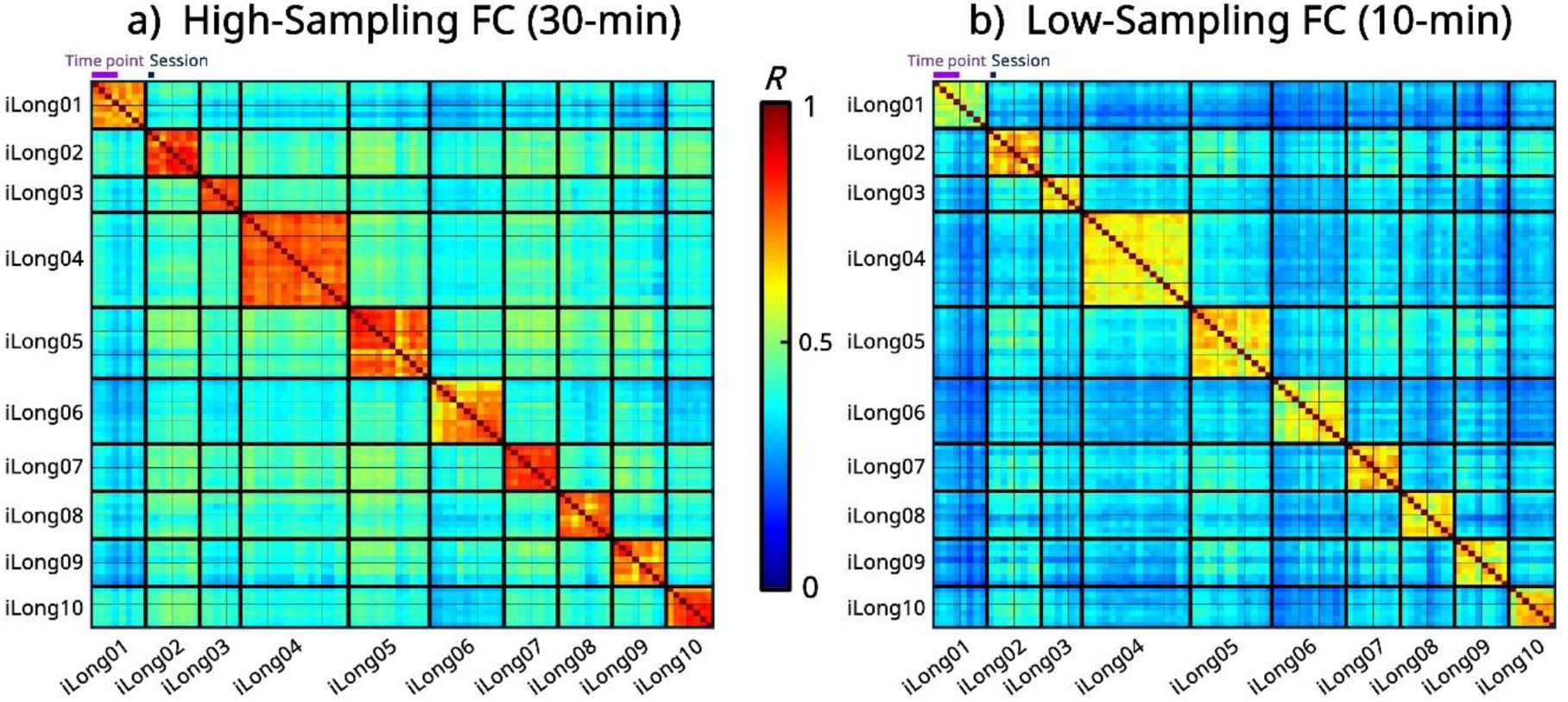
Extended per-participant scanning reveals robust temporal stability. **a)** We compared functional connectivity (FC) between cortical regions of interest (Seitzman300). Each cell of the matrix represents the similarity of FC between pairs of sessions (data amount matched to 30 minutes per session). The matrix shows high correlations within participants (indicated by warmer colors along the diagonal) and low correlations between participants (cooler colors), supporting that FC is individual-specific. Additionally, there were notable high correlations between time points within participants, as indicated by the similarity of colors across time points for each participant. **b)** In contrast to the left panel, this matrix was generated using the first 10 minutes of data after censoring high-motion frames, which is often the case in many consortium resting-state fMRI datasets. Correlations within participants were still notably higher than those between participants, but the mean within-participant similarity was significantly smaller than the left. Sufficient per-participant data—more than 30 minutes per session—would likely be needed to demonstrate that temporal stability can reach levels above 0.75.

### 1.4. Hierarchical network organization of functional stability

The vertex-level analyses (**Figs. 2-4**) showed that the majority of vertices and large-scale networks exhibited temporal stability. However, relatively lower stability was observed within sensorimotor and visual areas. To quantify whether specific networks differed in their longitudinal changes, we ranked networks from highest to lowest stability on average across the Standard Cohort participants (**Fig. 6a**). When stability estimates were averaged across networks within each category, association networks (default mode (DMN), frontoparietal (FP), cingulo-opercular action-mode (CO/AMN), dorsal attention (DAN), and salience (SAL)) showed significantly greater temporal stability than sensorimotor networks (visual (Vis), auditory (Aud), somatomotor ventral (SMv), and somatomotor dorsal (SMd); paired t-test, *t*(9) = 4.85, *p* < 0.001, Cohen’s *d* = 1.53; **Fig. 6b**). Note that although medial temporal lobe (MTL) and temporal pole (Tpole) are also association networks, they mostly overlap with low SNR regions and were thus considered separately (gray bars in **Fig. 6a**). This ranking was highly similar to the ranking of network similarity of the Prolonged Longitudinal Cohort (**Supp. Fig. 4**).

**Figure 6.**
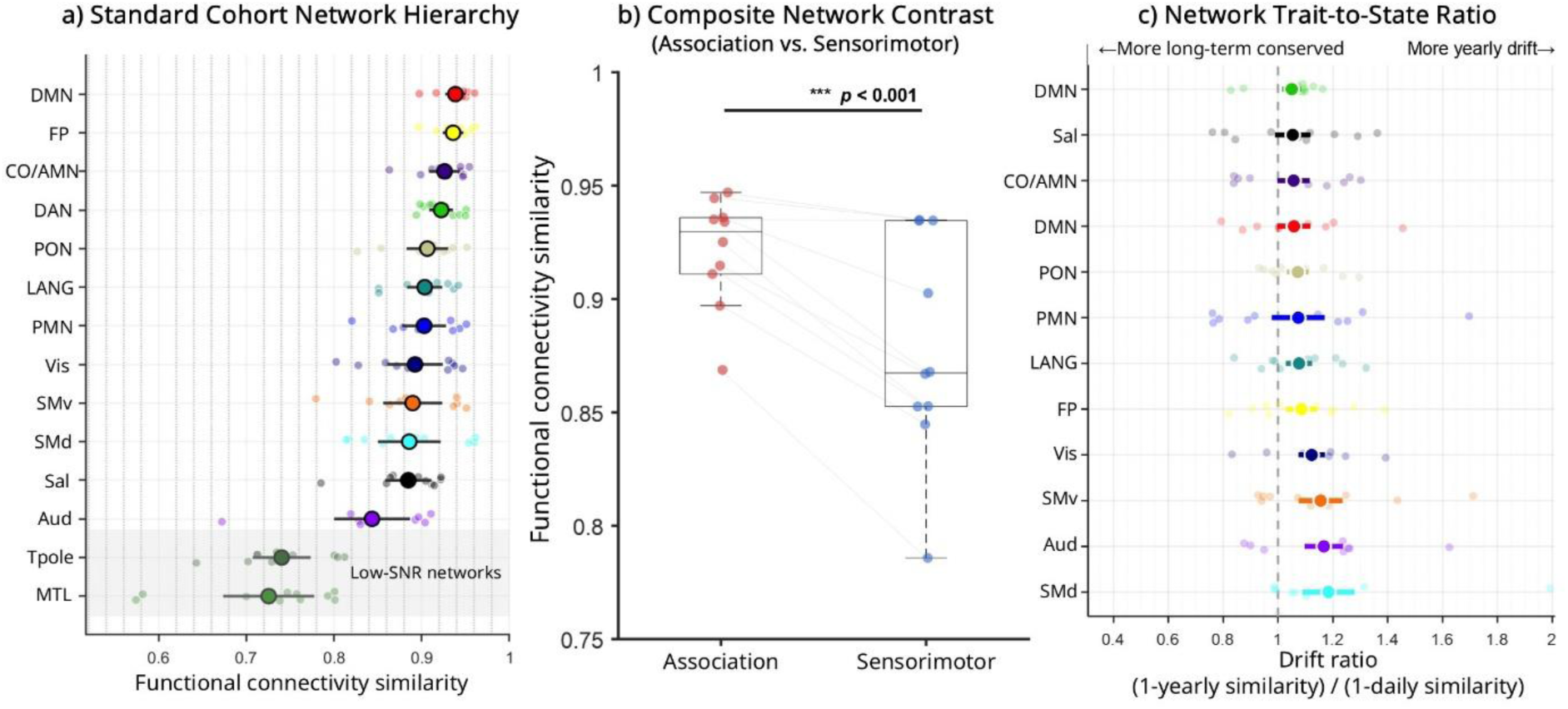
Identifying functional networks that are more stable across years. **a)** Network stability rank ordering of the Standard Longitudinal Cohort. The circles indicate the mean functional connectivity similarity for each network across participants with the solid lines showing its 95% confidence interval (individual points are from each participant). The means were ranked from most stable to least stable. **b)** Comparing mean stability between association and sensorimotor networks showed that association networks exhibited significantly greater temporal stability (each dot indicates one Standard Cohort participant). **c)** For each network, functional connectivity similarity was computed across sessions within the same time point (weekly stability) and across longitudinal time points (yearly stability). We computed a drift ratio (1-yearly similarity)/(1-daily similarity). Values near 1 indicate comparable divergence across timescales, whereas ratios >1 indicate greater long-term drift relative to short-term session variability. On average, networks show a drift ratio close to, but slightly exceeding 1, indicating subtle accumulating yearly drift in brain networks. This pattern was stronger in sensorimotor than association networks. (DMN, default mode network; FP, frontoparietal network; CO/AMN, cingulo-opercular action-mode network; DAN, dorsal attention network; PON, parieto-occipital network; LANG, language network; PMN, medial parietal network; Vis, visual network; SMv, somatomotor ventral; SMd, somatomotor dorsal; Sal, salience network; Aud, auditory network; Tpole, temporal pole; MTL, medial temporal lobe)

We next asked whether the pattern of functional network changes across years exceeded the amount expected from daily variability. To this end, we compared long-term divergence (1 - functional connectivity similarity across years) to short-term variability (1 - functional connectivity similarity across weeks; **Fig. 6c**). Across networks, these drift ratios were generally close to 1, with a modest tendency to exceed it, indicating that yearly and weekly variability are close to equivalent across networks with slightly larger changes seen across years. While generally similar, the largest yearly drift ratios were seen for sensorimotor networks, while the most stability was seen in association networks. In summary, all networks exhibited a profile of relatively high stability across years with only subtle accumulating drift across years, while the highest yearly stability was found in association networks.

### 1.5 Idiosyncratic network variants show longitudinal stability

While we have found that brain networks, as a whole, are longitudinally stable, we further assessed the stability of locations with the most person-specific features. Network variants are locations where an individual’s functional connectivity patterns differ the most from the group average (Seitzman et al., 2019). In past work, we have demonstrated that these locations are reliable across sessions over approximately a month (Dworetsky et al., 2024; Seitzman et al., 2019). Here we further examined how consistent these locations are over years, during which substantial neural and environmental changes accumulate, in order to examine the extent to which brain network variants could function as trait-like biological markers.

Consistent with our other measures, network variants within individuals demonstrated stability over time (**Fig. 7**). In the three Prolonged Cohort participants (prolonged years highlighted in pink), the within-participant Dice coefficient of the variants (*M* = 0.51, *SD* = 0.13) was significantly higher than the between-participant Dice coefficient (*M* = 0.18, *SD* = 0.02), *p* < 0.001 (permutation testing via label shuffling). This finding was replicated in the MyConnectome dataset and its 10-year follow-up (Dice = 0.5; see **Supp. Fig. 5** for the full timeline and brain maps).

**Figure 7.**
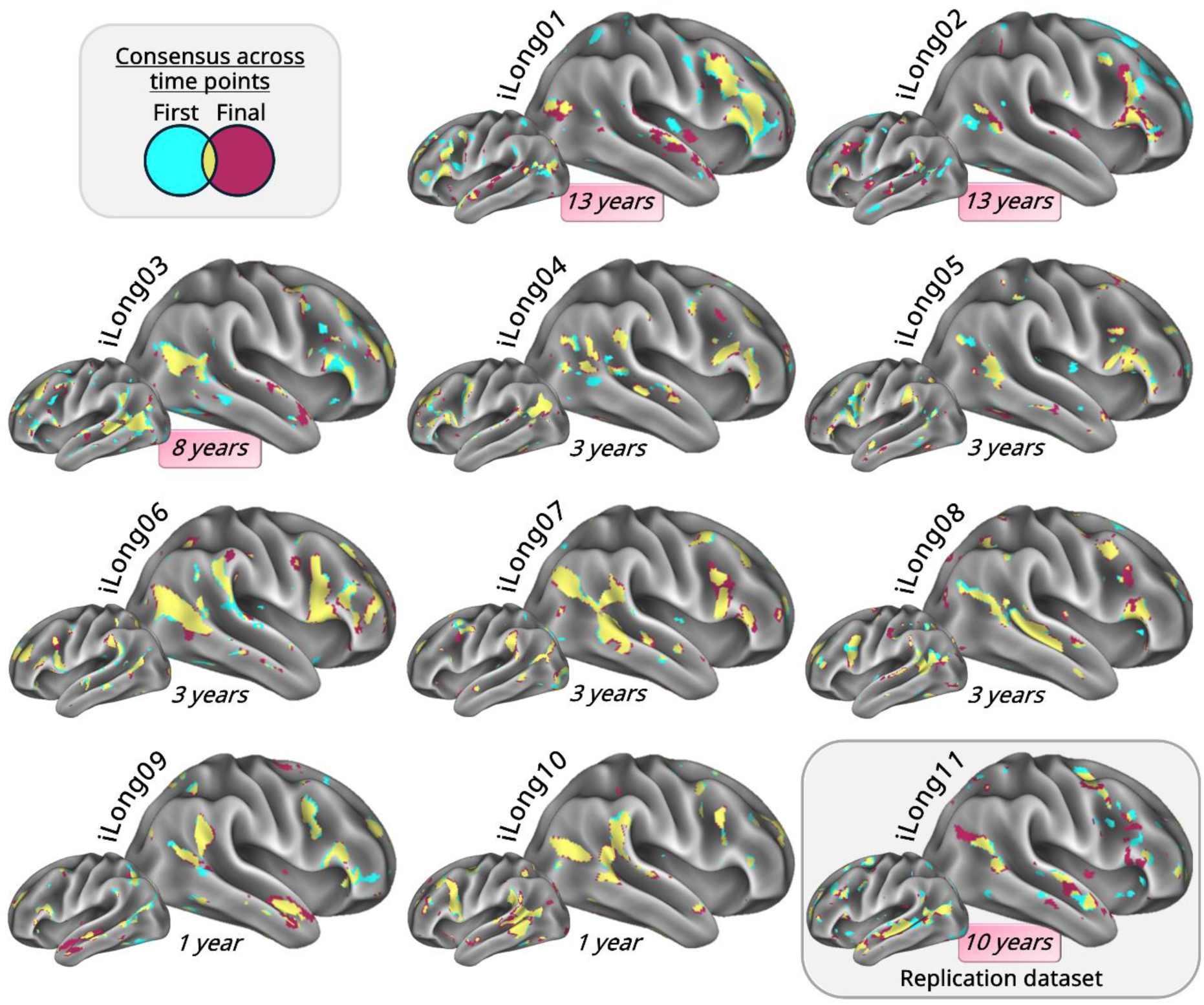
Idiosyncratic network variants overlap across time. Network variants were mapped for each participant at each time point. In line with prior studies, the location, topography, and size of variants varied across participants, consistent with their characterization as idiosyncratic features (Dworetsky et al., 2024; Seitzman et al., 2019). Notably, variants showed high overlap between time points within participants (marked in yellow), even across time spans of more than a decade. See also **Supp. Fig. 7** for the unthresholded variant maps and **Supp. Fig. 8** for the detailed composition of these overlapping variants, illustrating the network affiliation of each vertex.

The consensus was also notable in the Standard Cohort, with a mean Dice coefficient of 0.76 (*SD* = 0.05). When compared at a session-level (with matched, but more limited data of 30 minutes per session), we found that network variants overlapped similarly across yearly (*Dice M* = 0.58) and weekly timescales (*Dice M* = 0.60), where yearly comparisons significantly exceeded comparisons between participants (*Dice M* = 0.16; permutation *p =* 0.002; see **Supp. Fig. 6**). These findings confirm that even the most idiosyncratic features of functional brain organization are largely stable across years.

## DISCUSSION

Here, we showed that functional network organization is largely stable within individuals, even across extended time windows of over a decade. The brain’s stability and individual-specific nature indicate that it can encode functional characteristics of personhood that remain constant amid temporal fluctuations and life transitions. These observations refine current views of brain resilience and plasticity by highlighting the capacity of homeostatic mechanisms to maintain idiosyncratic network features. These insights also pave the way for a more comprehensive understanding of the neural mechanisms behind stability to pinpoint critical markers of healthy brain function for clinical applications.

### Functional networks as stable neural signatures of individual identity

Our findings suggest that functional networks putatively serve as trait-like markers, preserving stable characteristics in the face of changing life experiences. Previous studies demonstrated that resting-state functional connectivity is preserved across distinct mental (Cole et al., 2014; Du et al., 2025b; Gratton et al., 2018) and physiological states, including light sleep and anesthesia (Horovitz et al., 2009; Larson-Prior et al., 2011; Laumann & Snyder, 2021). We show that this stability extends even across prolonged periods during which participants experienced major life stressors and transitions, highlighting that network organization is robust not only to state fluctuations but also to long-term changes in life circumstances.

Our findings are also consistent with prior longitudinal reports using comparatively limited data (Badhwar et al., 2020; Horien et al., 2019; Jalbrzikowski et al., 2020). For example, individuals could be re-identified with comparable accuracy when scans were conducted on the same day versus when they were conducted 1.5 years apart (Jalbrzikowski et al., 2020). A study in mice has shown that mouse functional connectivity remained relatively stable from juvenile to adult stages across months once major developmental milestones had occurred (Millevert et al., 2025).

This stability is consistent with models proposing that functional connectivity reflects a statistical history of co-activations between brain regions (Petersen & Sporns, 2015; Poldrack et al., 2015), constrained by a combination of direct anatomical connections (Honey et al., 2009; Laumann & Snyder, 2021) and indirect network dynamics (Honey et al., 2009; Park & Friston, 2013). While some have proposed that these network dynamics may be highly malleable, the robust stability we find across more than a decade suggests that functional network dynamics are stably maintained to withstand contextual stressors, likely through homeostatic mechanisms (Laumann & Snyder, 2021). Thus, while more transient dynamics likely also exist (Calhoun et al., 2014; J. R. Cohen, 2018; Gonzalez-Castillo & Bandettini, 2018; Hutchison et al., 2013; Lurie et al., 2020), they are embedded within a dominant, stable network structure.

These stable features indicate that network organization may be a biological substrate of individual identity and complex cognition, characterizing our personhood. Differences in the spatial arrangement of functional regions have been strongly linked to inter-individual behavioral differences (Bijsterbosch et al., 2018; Finn et al., 2015; Miranda-Dominguez et al., 2014; Pierce et al., 2024). While such topographical variation is highly stable within individuals, it constitutes the biggest source of variation across individuals (Bijsterbosch et al., 2018). In accordance with this, topographical variation can predict inter-individual variation in higher-order cognitive abilities, such as fluid intelligence (Kong et al., 2019), that shape complex human behavior.

Taken together, the convergence of temporal stability, biological grounding, and cognitive relevance suggests that network features capture individually distinctive and enduring aspects of brain organization, constituting possible biological markers of individual identity.

### Reconciling stability with neural plasticity

Our data showing high stability in functional network topography across even a decade may, at first, appear surprising given findings in the literature on brain plasticity. However, we suggest these findings can be reconciled by considering that functional topography may be most malleable and subject to dramatic and lasting changes during critical developmental periods, after which network changes are largely focal (Gratton & Braga, 2026). This approach to plasticity would allow brain organization to remain stable and resilient to outside forces, supporting individual identity and protecting the brain from extreme changes at the whims of recent experiences.

Although not targeted in our study, additional sources of potential variation in brain networks are neurodevelopmental (Fair et al., 2009; Fransson et al., 2007; Gao et al., 2015; Gratton et al., 2018; Grayson & Fair, 2017; Nielsen et al., 2019; Satterthwaite et al., 2012; Smyser et al., 2010) and aging-related processes (Campbell et al., 2012; Madden et al., 2010; Spreng & Schacter, 2012). In the developing brain, networks become more specialized, with within-network connectivity strengthening as they mature into more cohesive and functionally distinct systems (Baum et al., 2017; Gao et al., 2015; Grayson & Fair, 2017), and potentially more individualized into adulthood (Demeter et al., 2025). In contrast to these common developmental patterns, patients with congenital or perinatal damage can develop atypical large-scale network organization, as evidenced by the unusual motor organization in patient PS1 (Laumann et al., 2021) and language network in patient EG (Tuckute et al., 2022) (see also (Newport et al., 2022)). In addition, extreme variation in sensory or motor experience can lead to idiosyncratic variation in functional relationships of primary sensory and motor regions (Amaral et al., 2024; Schone et al., 2025). These findings are consistent with animal literature, demonstrating that, while proto-functional organization is present from birth (Arcaro & Livingstone, 2017), sensory deprivation during critical periods can lead to substantial reorganization in functional responses (Arcaro et al., 2017).

However, after critical development, we show that brain organization remains quite stable in adulthood. This is consistent with prior literature: in Patient EG, who exhibits atypical language organization due to congenital absence of the left temporal lobe, the language network remained stable during the 3-year longitudinal observation period (Tuckute et al., 2022). Similarly, the functional connectivity of primary visual cortex (V1) of people with congenital blindness has been shown to be stable over time and contexts, retaining its idiosyncratic profile (Amaral et al., 2024).

Even in the face of dramatic changes in sensory experience, reorganization appears unlikely in adulthood, as longitudinal neuroimaging of three adults before and after arm amputation showed that despite the loss of peripheral input, the intrinsic organization of hand-related activity in the sensorimotor cortex persisted, resisting remapping (Schone et al., 2025). Rare departures have been reported primarily under extreme cases, such as major medical interventions (Mitchell et al., 2020; Preller et al., 2020; Seitzman, Anandarajah, et al., 2023; Seitzman, Reynoso, et al., 2023).

As we have demonstrated, functional network architecture may be largely fixed in adulthood, consistent with these case reports. This stability is particularly notable when considering major life events that occurred in our participants, including socio-cultural and health phenomena (e.g., the COVID-19 pandemic), as well as personal experiences (e.g., childbirth and rearing, job transitions, cross-country moves, and death of a parent). These examples align with events listed in the Social Readjustment Rating Scale (Holmes & Rahe, 1967), which identifies life-altering occurrences that are significant stressors impacting health and well-being.

When network modulations take place in response to experience or symptoms, they are likely to take a more subtle and transient form, affecting correlation strength among specific relevant connections. One longitudinal study that scanned participants repeatedly before, during, and after wearing arm casts showed fluctuations in the magnitude of functional connectivity, but these were focal to primary motor regions and reversed with removal of the casts (Newbold et al., 2020). Lynch and colleague (Lynch et al., 2024) found evidence of an enlarged salience network in individuals with depression with a topography that did not change over time; however the magnitude of specific correlations was altered with changes in symptoms. Over time, these more focal changes may accumulate to longer lasting variation (perhaps associated with the slight drift measured in our study in **Fig. 6**), but this process may extend over decades in adulthood. This functional stability may provide a substrate for our enduring stability of self and resilience to outside forces.

Identifying when idiosyncratic features first emerge in the developing brain and the temporal extent of their persistence throughout aging represents an important direction for future investigations. Our findings of stability set the stage for such work by delineating a temporal regime in young, healthy adults in which network measures are predominantly stable.

### Clinical implications of stable functional networks

Our findings underscore the value of precision fMRI in disentangling stable trait-like neural features from more transient state-dependent fluctuations. Such stable individual baselines may be crucial for identifying subtle or progressive changes associated with pathology. Distinguishing between stable functional topography and time-varying connectivity may enable early identification of neurodegenerative disease progression, treatment effects, and targets for neuromodulation, along with understanding the impact of life events on mental health.

As an example, Lynch and colleagues (Lynch et al., 2024) showed that the topological features of the salience network, including its shape and size, remained stable over months, and could even be observed prior to the onset of depression in children. Interestingly, more fine-scale fluctuations within this variant network could be used to predict fluctuations in anhedonia and anxiety symptoms over time. This exemplifies how functional networks may serve as trait-like markers of risk for clinical symptoms and allow for sensitive detection of subtle markers linked to changing brain health.

### Sources and limits of temporal stability

Although we demonstrated that functional brain organization was predominantly stable over time, some variability existed that can be linked to measurement issues. For example, lower stability was observed in the inferior and medial temporal lobe and orbitofrontal cortex, where the BOLD signal is more difficult to estimate due to magnetic field inhomogeneities and physiological noise (Bianciardi et al., 2009; Ojemann et al., 1997b; Thomas Yeo et al., 2011). Relatively lower stability was also observed in the Prolonged Cohort which may have been associated in part with differences in scanner types and sequences (although these may also be linked to the longer inter-scan intervals in this group).

Another potential factor that can cause apparent changes in functional connectivity is data quantity. Compared to 30 minutes, using 10 minutes of resting-state fMRI data resulted in dramatically lower stability across sessions and time points (**Fig. 5**), due to sampling variability in MRI functional connectivity measures (Laumann et al., 2015). This aligns with past literature observing that shorter scan durations resulted in poorer reliability of functional connectivity (Noble et al., 2019). Reducing measurement noise in functional connectivity is particularly important in longitudinal studies, as biases induced by noise may otherwise look like meaningful changes (e.g., if people move more with age, leading to added data loss, apparent longitudinal effects of aging may be driven by sampling variability). Therefore, it is critical to acquire precision-level data and adopt advanced denoising techniques to robustly detect longitudinal changes (Vinci-Booher et al., 2025). Moreover, while it is uncommon in many longitudinal studies, having multiple sessions at each time point, as in our dataset, can be highly useful to separate longitudinal changes from day-to-day variability.

We also observed relatively larger changes in visual and motor regions (**Fig. 6**), consistent with prior work examining daily variability within individuals in functional networks (Gratton et al., 2018; Laumann et al., 2015). Analyses of the initial MyConnectome dataset showed that session-to-session variability was greatest in visual and somatomotor regions over a year, whereas relatively less variability was observed in the frontoparietal, default mode, ventral attention, and medial parietal regions (Laumann et al., 2015). These fluctuations were interpreted as reflecting state-related factors, particularly variations in arousal (Tagliazucchi & Laufs, 2014), given that they also tracked with caffeination status. Recent work has shown that the strongest brain-wide associations with socioeconomic factors are concentrated in primary motor and sensory networks in children and exhibit strong spatial similarity to arousal-related patterns, including sleep duration, norepinephrine signaling, and stimulant effects (Marek et al., 2026). These findings suggest that sensorimotor networks are particularly sensitive to fluctuations in arousal state, sleep, and other environmental exposures. Because these influences can vary substantially within individuals over time, sensorimotor networks may be especially susceptible to longitudinal changes, and consequently, exhibit lower stability than higher-order association networks that can be relatively insulated from state-dependent fluctuations. Interestingly, in our data these patterns of daily variability in sensorimotor regions were mirrored, and indeed slightly amplified across years, suggesting that they may accumulate drift over time. In contrast, in this age group of healthy adults, association regions of the brain were relatively more stable, exhibiting trait-like tracking of person-specific networks even over long time windows.

## CONCLUSION

We found that functional brain networks within individuals were preserved across extended timescales of over a decade despite ongoing neural dynamics. When scanners and protocols were matched, we observed minimal changes over a 2-year period, only subtly exceeding the variability seen in brain networks from day to day. These findings demonstrate that functional networks track individual identity and can withstand changes in the face of major life experiences.

## METHODS

### Datasets

Our participants consist of 10 young, healthy adults (five females and five males, aged 23-30 years, with a mean age of 26, at the time of the first scan). In each participant, we collected structural and functional MRI measures (≥60 minutes of resting-state fMRI per time point) across two or three time points separated by up to 13 years (see *Data Acquisition*).

We divided the participants into two cohorts: the **Prolonged Longitudinal Cohort** and the **Standard Longitudinal Cohort** (**Fig. 1**). Three participants from the Prolonged Longitudinal Cohort (iLong01-03) were also included in the Standard Longitudinal Cohort (iLong01-10), but we distinguished these cohorts to highlight that Prolonged Cohort sessions were conducted with extended delays (8 or 13 years) using different scanners and imaging protocols (see *Table 1*). Standard Cohort sessions were conducted with shorter intervals (1-3 years; *Table 3*) using the same scanner and protocols. We replicated our Prolonged Cohort findings using the publicly available MyConnectome data (male, 45 years old at the time of the first scan) supplemented by additional resting-state fMRI data collected a decade later (iLong11; *Table 2*).

**Table 1.** Participant demographics and scanning parameters for the Prolonged Longitudinal Cohort. The Prolonged Cohort consists of three participants. Two participants’ data span approximately 13 years, from 2009 to 2022, and one participant’s data span 8 years, from 2014 to 2022. Participant demographic information includes age at time point (TP) 1 and sex. The number of fMRI sessions collected per TP, the number of runs per session, repetition time (TR; in seconds), and scan duration (in minutes (Mins)) are also shown. The minimum scan duration is 60 minutes (58 minutes after processing and censoring high head motion frames), which is sufficient for reliable examination of individual cortical network maps (Gordon et al., 2017). At the second time point (2022), data were collected using different scan parameters (refer to Table 3).

|  | ID | Age | Sex | TP1 | No. Session /TP1 | No. Run /Session | TR | Mins/TP1 | TP2 |
| --- | --- | --- | --- | --- | --- | --- | --- | --- | --- |
| **Prolonged** | iLong01 | 23 | F | 2009 | 1 | 6 | 1.375 | 60 | 2022 |
|  | iLong02 | 23 | M | 2009 | 1 | 6 | 1.375 | 60 | 2022 |
|  | iLong03 | 24 | F | 2014 | 10 | 1 | 2.2 | 300 | 2022 |

**Table 2.** Participant demographics and scanning parameters for the replication dataset. From the initial MyConnectome dataset (100 sessions), we selected six sessions collected within a two-month period (10/29/2012-12/10/2012), and for the second time point in 2022, we used all available sessions acquired within a comparable two-month window (10/03/2022-11/19/2022). The first four sessions at this second time point were collected with a multiecho sequence and the remaining six with a single-echo sequence. To combine these datasets, we used the second echo from the multiecho scans, which had an echo time closest to the single-echo acquisition. Other acquisition parameters (e.g., TR, reported in seconds, and voxel size) also differed; see *Data Acquisition* below for details.

|  | ID | Age | Sex | TP1 | TP2 | No. Session /TP1 | No. Session /TP2 | No. Run /Session | TR 1 | TR 2 | Mins /TP1 | Mins /TP2 |
| --- | --- | --- | --- | --- | --- | --- | --- | --- | --- | --- | --- | --- |
| **Prolonged** | iLong11 | 45 | M | 2012 | 2022 | 6 | 3 | 1~4 | 1.16 | 1.49, 1.25 | 61 | 64 |

**Table 3.** Participant demographics and scanning parameters for the Standard Longitudinal Cohort. The controlled cohort, using matching scanner and scan protocols, consists of 10 participants. All scans were conducted at Northwestern University. Participants underwent either two or three scan time points. The minimum scan duration per time point is 89 minutes. iLong01, iLong02, and iLong03 are part of the Prolonged Cohort, with the second time point (2022) of the Standard Cohort for these participants corresponding to the second time point of the Prolonged Longitudinal Cohort.

|  | ID | Age | Sex | No. TP | TP1 | TP2 | TP3 | No. Session /TP1 | No. Session /TP2 | No. Session /TP3 | No. Run /Session | TR | Mins /TP1 | Mins /TP2 | Mins /TP3 |
| --- | --- | --- | --- | --- | --- | --- | --- | --- | --- | --- | --- | --- | --- | --- | --- |
| Standard | iLong01 | 35 | F | 2 | 2021 | 2022 | - | 4 | 4 | - | 9~11 | 1.1 | 218 | 198 | - |
|  | iLong02 | 34 | M | 2 | 2021 | 2022 | - | 4 | 4 | - | 9~11 | 1.1 | 213 | 193 | - |
|  | iLong03 | 29 | F | 2 | 2020 | 2022 | - | 4 | 2 | - | 7~10 | 1.1 | 153 | 89 | - |
|  | iLong04 | 29 | M | 3 | 2019 | 2020 | 2022 | 5 | 8 | 4 | 6~10 | 1.1 | 189 | 342 | 178 |
|  | iLong05 | 26 | M | 3 | 2019 | 2021 | 2022 | 4 | 4 | 4 | 7~11 | 1.1 | 153 | 198 | 168 |
|  | iLong06 | 27 | M | 3 | 2019 | 2021 | 2022 | 4 | 4 | 4 | 6~9 | 1.1 | 158 | 149 | 178 |
|  | iLong07 | 30 | F | 2 | 2019 | 2022 | - | 4 | 4 | - | 7~9 | 1.1 | 163 | 173 | - |
|  | iLong08 | 26 | F | 2 | 2019 | 2022 | - | 4 | 4 | - | 7~10 | 1.1 | 153 | 183 | - |
|  | iLong09 | 28 | M | 2 | 2021 | 2022 | - | 4 | 4 | - | 7~10 | 1.1 | 144 | 178 | - |
|  | iLong10 | 26 | F | 2 | 2021 | 2022 | - | 4 | 4 | - | 7~10 | 1.1 | 153 | 183 | - |

#### Prolonged Longitudinal Cohort

Two participants from the Prolonged Longitudinal Cohort (iLong01 and iLong02) were scanned at the University of California, Berkeley, in 2009 (IRB: 2010-02-781) for their first time point. A single session, consisting of 60 minutes of resting-state fMRI data, was collected from each participant. These participants were also included as healthy controls in previous studies (Gratton et al., 2012; Nomura et al., 2010).

The third participant (iLong03) was scanned at Washington University in St. Louis in 2014 (IRB: 201402100) for their first time point and is part of the Midnight Scan Club data (Gordon et al., 2017). Ten fMRI sessions across approximately three weeks were collected from this participant, accumulating 300 minutes of resting-state fMRI data.

Subsequent time points followed the data collection procedures described in the Standard Longitudinal Cohort.

#### Replication dataset

The replication participant (iLong11) was scanned at University of Texas at Austin in 2012, with the data released in 2014 as the MyConnectome dataset (Laumann et al., 2015; Poldrack et al., 2015)(IRB determined study to be not human subject research). This dataset was supplemented with additional longitudinal data collected in 2022 at Stanford University.

We began by investigating the temporal stability of functional brain organization in the Prolonged Cohort to address whether brain organization remains stable over extended time windows. We assessed temporal stability in (i) network assignment across the whole brain, (ii) spatial correlation of the functional connectivity seed map of each vertex, and (iii) idiosyncratic network variants. These measures represent systematic organization of intrinsic functional connectivity that can capture individual-specific features.

#### Standard Longitudinal Cohort

We then conducted more controlled investigations of the functional organization measures by collecting data in 10 participants with matched scan protocols. The scans were conducted between 2019 and 2022 at two or three longitudinal time points at Northwestern University (IRB: STU00208793) using the same scan protocols. For these time points, participants completed four sessions of fMRI data collection across different days (with three exceptions: iLong03 completed two sessions at time point 2, and iLong04 completed five and eight sessions at time points 1 and 2, respectively). This allowed us to collect more than 89 minutes of resting-state data at each time point.

### Data Acquisition

All datasets included structural and functional MRI. Functional MRI measures included resting-state scans, which were the primary focus of analysis in this study. In these resting-state fMRI scans, participants were instructed to remain relaxed, still, and keep their eyes open, focusing on a fixation point. For each institution, scanner make and scanning parameters were as follows:

- University of California Berkeley: Imaging was conducted using a 3T Siemens MAGNETOM Trio, equipped with a 12-channel head coil. Structural images were acquired using an axial MP-RAGE 3D T1-weighted sequence (repetition time (TR) = 2,300 ms, echo time (TE) = 2.98 ms, flip angle = 9°, isotropic voxel size = 1 mm^3^). A gradient-echo EPI BOLD sequence was applied with the following parameters: TR = 1,370 ms, TE = 50 ms, voxel size = 2.34375 x 2.34375 x 3.85 mm, and twenty-four 3.85-mm-thick axial slices. See (Gratton et al., 2012; Nomura et al., 2010) for further details.
- Washington University in St. Louis: Imaging was carried out on a 3T Siemens MAGNETOM Trio. Structural MRI was conducted across two days, collecting T1-weighted images (TR = 2,400 ms, TE = 3.74 ms, inversion time (TI) = 1,000 ms, flip angle = 8°, isotropic voxel size = 0.8 mm^3^, 224 slices), T2-weighted images, MRAs, and MRVs. A gradient-echo EPI BOLD sequence was used with these settings: TR = 2,200 ms, TE = 27 ms, flip angle = 90°, isotropic voxel size = 4 mm^3^, and 36 axial slices. Refer to (Gordon et al., 2017) for further information.
- Northwestern University: Data were collected using a 3T Siemens MAGNETOM Prisma scanner, equipped with a 64-channel head coil. Structural images were collected using 3D T1-weighted images (TR = 2,300 ms, TE = 1.9 ms, TI = 1,180 ms, flip angle = 7°, isotropic voxel size = 0.8 mm^3^, 176 slices). A gradient-echo EPI BOLD sequence was employed with the following parameters: TR = 1,100 ms, TE = 25 ms, flip angle = 80°, isotropic voxel size = 2.5 mm^3^, and 60 axial slices. Multiband imaging was used with an acceleration factor of 4.
- University of Texas at Austin: Imaging was conducted using a 3T Siemens Skyra MRI scanner with a 32-channel head coil. Of the 100 sessions the initial MyConnectome dataset provides, we selected sessions 15, 18, 21, 24, 27, and 30 that were high-quality sessions in which at least 91% of frames were retained after censoring high-motion frames based on filtered framewise displacement (fFD) calculations to match the data quantity available in the post-10-year protocols. These sessions also included complete anatomical, field map, and functional data. T1-weighted images were acquired using an MP-RAGE sequence (TR = 2,400 ms, TE = 2.14 ms, TI = 1,000 ms, flip angle = 8°, GRAPPA factor = 2, isotropic voxel size = 0.8 mm^3^, 256 slices). Resting-state fMRI data were acquired using a multiband EPI sequence (TR = 1,160 ms, TE = 30 ms, flip angle = 63°, voxel size = 2.4 × 2.4 × 2 mm, 68 slices, multiband factor = 4). Starting with session 27, the slice count was reduced to 64 due to an update that altered the minimum TR. Acoustic noise reduction was attempted using an Optoacoustics active noise-cancellation system. Refer to (Poldrack et al., 2015) and associated dataset documentation for further acquisition details.
- Stanford University: Imaging was performed at the Stanford Center for Cognitive and Neurobiological Imaging using a 3T GE SIGNA Premier scanner with a Nova 32-channel head coil. T1-weighted images were acquired using a 3D MP-RAGE sequence (TR = 3,000 ms, TE = 3.548 ms, TI = 1,060 ms, flip angle = 8°, isotropic voxel size = 0.8 mm^3^). The first four sessions employed a multiecho sequence for resting-state data (TR = 1,490 ms, TEs = 13.4, 36.4, and 59.4 ms, flip angle = 53°, isotropic voxel size = 2.8 mm^3^). From the 3 echoes, we selected the second echo, which had a TE closest to the single-echo scans. For the subsequent six sessions, data were acquired on the same scanner using a 48-channel head coil with a single-echo EPI sequence. (TR = 1,250 ms, TE = 30 ms, flip angle = 53°, 2.2 mm isotropic voxels, 64 slices, multiband factor = 4).

### Data and Code Availability

Not all data used in this study can be made publicly available due to restrictions in the original IRB protocols. However, a subset of the data is accessible at OpenNeuro (Midnight Scan Club (MSC) dataset https://openneuro.org/datasets/ds000224/versions/00001) and NIMH’s NDA (iNetworks dataset); only the data from participants who consented to share with the NDA during their final questionnaire is available on the NDA platform.

MyConnectome and the follow-up data are publicly available at OpenNeuro (https://openneuro.org/datasets/ds000031/versions/2.0.2).

All data processing and analysis code (custom Matlab scripts, https://github.com/GrattonLab/iLongitudinal) are publicly available.

### Preprocessing

We used fMRIPrep 23.2.0 to preprocess data, with consistent structural and functional workflows across cohorts, except for a few site-specific differences detailed below. Structural T1-weighted (T1w) images underwent intensity non-uniformity correction with N4BiasFieldCorrection, skull stripping using Advanced Normalization Tools (ANTs) with the OASIS30ANTs template, brain tissue segmentation into gray matter, white matter, and cerebrospinal fluid using fast FSL, and surface reconstruction with FreeSurfer 7.3.2 (recon-all). The brain mask was refined using a Mindboggle-based method that reconciles ANTs- and FreeSurfer-derived segmentations, producing a hybrid brain mask to improve cortical gray matter delineation. Volume-based spatial normalization to MNI152NLin6Asym and MNI152NLin2009cAsym space was performed via nonlinear registration using ANTs.

For functional preprocessing, a reference volume was generated from the BOLD time series following susceptibility distortion correction, where applicable. For data (aside for those acquired at the University of California, Berkeley) a dual-echo Gradient-Recalled Echo (GRE) sequence was used to estimate the B_0_ fieldmap.

Slice timing correction was then performed only for data collected at Washington University in Saint Louis, given its longer TR and single band acquisition. Head motion correction was performed using mcflirt (FSL), estimating rigid-body motion parameters (translations and rotations), along with the corresponding transformation matrices, prior to any spatiotemporal filtering. The BOLD reference was then co-registered to the T1-weighted anatomical image using bbregister (FreeSurfer), employing boundary-based registration with six degrees of freedom. All spatial transformations—including motion correction, susceptibility distortion correction, and co-registration—were combined and applied in a single interpolation step using nitransforms with cubic B-spline interpolation.

### Functional Connectivity Processing

Resting-state fMRI data underwent additional denoising procedures following the pipeline described by Power et al. (2014), to reduce noise from physiological sources of non-interest such as respiration, heart rate, and head motion. BOLD time series were intensity-normalized using mode-1000 normalization. The time series were then demeaned and detrended to remove slow scanner-related fluctuations.

Nuisance regression was performed to remove motion (Friston 36-parameter model with white matter, CSF, and global signals, along with their squares and temporal derivatives). High-motion frames were censored using framewise displacement (FD), based on filtered FD (fFD>0.1 mm), where a low-pass filter (<0.1 Hz) was applied to the motion parameters to attenuate high-frequency respiration-related artifacts (Gratton et al., 2020). Additionally, the first five frames of each run and any segments shorter than five frames after censoring were also marked for removal. Censored frames were interpolated over using power-spectral matched interpolation. After interpolation, a temporal bandpass filter (0.009-0.08 Hz) was applied to the data. A final round of demeaning and detrending was then performed. The resulting cleaned BOLD data were used for subsequent functional connectivity analyses, with censored frames excluded based on the created temporal masks.

### Volume-To-Surface Mapping

After preprocessing, BOLD timeseries data were mapped to each participant’s native cortical surface. This native surface was then aligned to the FSLR-32k surface following Gordon et al.’s (2016) procedure. All subsequent data analyses were conducted on the cortical surface, except for the creation of functional connectivity similarity matrices of the cortical volume (**Fig. 5 and Supp. Figs. 2 and 3**), which were based on volumetric cortical regions of interest (‘Seitzman 300’; see below for details).

### Data Analysis

We discarded runs shorter than 2.75 minutes after censoring high-motion frames, resulting in the removal of one run each from iLong04 (4th session, 1st run), iLong06 (7th session, 2nd run), and iLong10 (2nd session, 6th run). Additionally, we discarded sessions shorter than 30.25 minutes after censoring, which led to the removal of one session from iLong04 (1st session), iLong06 (5th session) and iLong10 (2nd session). This was based on prior work demonstrating that approximately 30 minutes of low-motion data are required for highly reliable functional connectivity estimation (Gordon et al., 2017). For any session-level comparisons (**Figs. 3b, 4b-6; Supp. Figs. 2 and 6**), the data were matched to this duration to avoid confounding session differences with differences in data quantity.

When we censored low-SNR regions, we used a group-level binary map identifying vertices with low temporal signal-to-noise ratio (tSNR). This map excludes vertices with intensity-normalized signal values below 750 during functional connectivity processing.

#### Template-matching

We employed template-matching techniques to assign vertices in functional connectivity seed maps to networks. This method uses group-averaged network seed maps generated from the WashU-120 dataset (consisting of an independent set of 120 young adults; see Dworetsky et al., 2021 for details on template creation) to identify 14 distinct functional networks. The Dice coefficient was calculated between each of the 14 binarized network templates and each participant’s functional connectivity map at each vertex. The vertex was assigned to the network with the highest Dice coefficient using a winner-take-all approach. This process was applied to all of the vertices in the cortical surface data and repeated independently at each time point (or session, for some analyses) to assess network assignment consensus over time. For time-point-level maps, all available data were used to maximize individual-level precision (**Figs. 2a and 3a**), while for session-level maps, the data amounts were matched (30.25 minutes) to ensure consistency for statistical comparisons (**Fig. 3b**). Consensus maps were generated by assigning each vertex to the network with which it was consistent across time points or sessions.

#### Seed-based correlation mapping

As a second step, we examined the temporal stability of continuous vertex-wise functional connectivity using seed-based correlation analysis. We created surface-based functional connectivity seed maps by computing correlations between every pair of vertices at each time point. Then, to assess temporal consistency, we compared these vertex-level connectivity maps across two time points (or sessions) using Pearson correlation. **Figure 2b** presents the resulting spatial correlation maps for the Prolonged Cohort. **Figure. 2c** provides a dot plot of the values averaged across the cortex after censoring low tSNR vertices. **Figure 4** displays the results for the Standard Cohort. For comparisons across time points and participants in the **Figure 4a** matrix, data were matched to 86 minutes per time point, corresponding to the maximum amount of low-motion data available across all participants after censoring. In contrast, to maximize within-participant precision for the **Figure 4a** brain maps (which involved comparisons across time points within individuals but not across participants), data were retained at 136 per time point, except for iLong03, who had a maximum 86 minutes per time point. For session-level comparisons shown in **Figure 4b**, data amounts were matched to 30.25 minutes per session.

#### Volumetric ROI-based analysis of the similarity of functional connectivity maps

We also conducted analyses of the similarity in functional connectivity between volumetric regions of interest (ROI). We used previously defined atlas coordinates (‘Seitzman 300’; (Seitzman et al., 2020)) as a reference for the ROIs. After excluding censored frames with high head motion, the preprocessed BOLD time series for gray matter voxels were averaged for each ROI. The Pearson correlation coefficients were computed between the mean time series of every pair of ROIs, resulting in a 300 x 300 correlation matrix for each participant. This correlation matrix was created for each session and the upper triangle was compared across sessions and participants to generate a similarity functional connectivity matrix, as shown in **Figure 5**. This figure compares similarity matrices when using 30 minutes versus 10 minutes. **Supplementary Figure 2** presents results from comparisons of correlation matrices across sessions, shown as a dot plot using sessions from the furthest two time points. **Supplementary Figure 3** shows linear fixed-effect modeling results with these volume-based analyses.

#### Linear fixed-effects modeling of functional connectivity similarity

To quantify the extent to which elapsed time between sessions contributes to longitudinal functional connectivity similarity, we employed linear fixed-effects models on pairwise session comparisons using the volumetric Seitzman 300-ROI atlas (**Supp. Fig. 3**). For each participant, a 300x300 functional connectivity matrix and a corresponding voxel-wise tSNR map were generated for every session. ROIs were subjected to a two-tier censoring process: first, 11 ROIs with a group-mean tSNR below 30 were excluded. Second, for each participant, ROIs falling within the bottom 10^th^ percentile of that individual’s specific tSNR distribution were censored. For the remaining data, functional connectivity similarity was defined as the Pearson correlation between corresponding connectivity profiles of session pairs.

For each ROI, models were implemented using the *fitlm* function in Matlab with the following specification:

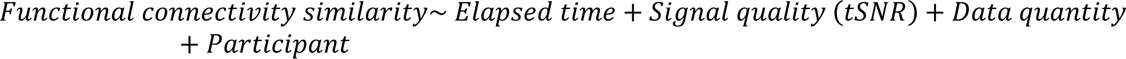

The inter-session interval (elapsed time in days) served as the primary fixed effect. We also included the mean tSNR of the session pair and the mean number of retained data points after censoring high-motion frames as nuisance covariates to control for signal quality and data quantity (ranged from 1,557-2,915 volumes), respectively. A ‘participant baseline’ variable was included to account for inter-individual differences in stability of networks. All predictors were standardized (z-scored) prior to model fitting to facilitate comparison across variables with different scales and to ensure that the intercept represented the expected similarity at the baseline. To assess the relative contribution of each predictor, variance partitioning was derived from the ANOVA tables obtained using Matlab’s *anova* function. For each factor, the effect size was quantified using *ω*^2^, calculated as follows (J. Cohen, 1973):

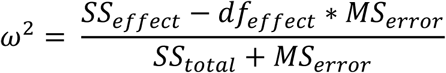

#### Network variants

We defined network variants as locations within an individual’s brain where the functional connectivity patterns differ significantly from the group average, following (Seitzman et al., 2019). For each participant, the functional connectivity map of every cortical vertex was correlated with the same vertex’s seed map of the reference group-average data (WashU-120; (Power et al., 2011)). This procedure produced an individual-to-group similarity map, where each vertex in the map reflects a spatial correlation value (**Supp. Fig. 7**). This similarity map was then thresholded to the lowest 10% to identify regions where the individual’s connectivity was least similar to the group. These regions were then binarized to define potential variant locations. Regions with low-SNR were excluded, using a vertex-wise map of low-signal areas, as described in Seitzman et al. (2019). The final binarized maps, created for each time point using all available data for each participant (**Supp. Fig. 5**), were matched with Dice coefficient to assess consistency across time (**Fig. 7**).

**Supplementary Figure 6** shows a dot plot of variant comparisons across the furthest two time points using the Dice coefficient, with matched data amounts (30.25 minutes per session). We also generated **Supplementary Figure 8** to examine the network variant composition and its temporal stability. We first generated network composition maps for each participant at each time point by combining binary variant maps (thresholded at the bottom 10% of spatial correlation values and SNR-masked) with the corresponding template-matching network topography. Each variant vertex was labeled according to its assigned network. To identify stable variants, we computed a consensus map for each participant, defined as vertices that consistently appeared as variants across all time points.

#### Permutation-based statistical tests

We conducted permutation tests for the Standard Longitudinal data to compare within-participant similarity across distant time points (red dots) against between-participant similarity (black dots). The same permutation procedure was applied to template matching consensus (**Fig. 3b**), functional connectivity similarity (**Supp. Fig. 2**), and network variant overlap (**Supp. Fig. 6**), with the specific comparisons tailored to each analysis. We employed group-level permutation testing (10,000 permutations), treating the participant as the primary unit of inference. For each participant, we calculated the mean difference in similarity between the two conditions (within versus between comparisons). We used the two temporally most distant time points for the within-participant measure. To generate a null distribution, we employed a label-shuffling approach, where the condition labels were randomly reassigned. The group-level *p*-value was determined by comparing the observed mean difference to this permutation-based null distribution.

For the Prolonged Longitudinal data, we conducted a non-parametric permutation test comparing three within-participant differences (one for each participant) to 24 between-participant differences (eight for each participant). Unlike the approach used for the Standard Longitudinal data, this pooled analysis of aggregating across participants was done to avoid low statistical power in testing whether within-participant consistency exceeds between-participant similarity. Within-participant consistency was quantified by comparing each participant’s two time points (e.g., iLong01 time point 1 versus time point 2), and between-participant similarity was calculated from all pairwise comparisons between different participants across all combinations of time points (e.g., iLong01 time point 1 versus iLong02 time point 2), yielding a total of 24 comparisons. The observed difference in mean between the within-participant group (n=3) and between-participant group (n=24) was compared against a null distribution generated from 10,000 random permutations of the group labels.

## ACKNOWLEDGEMENTS

This work was supported by NIH R01MH118370 (CG) and R01NS124738 (CG). Data for this work were stored and analyzed using HPC resources at the Research Computing Center at Florida State University, computational resources provided for the Quest high performance computing facility at Northwestern University, and the Illinois Campus Cluster, operated by the Illinois Campus Cluster Program (ICCP) in collaboration with the National Center for Supercomputing Applications (NCSA), with support from the University of Illinois Urbana-Champaign. We would like to thank Sunjae Shim and Jaime Rios for assistance with scanning in the MyConnectome follow-up study.

## Supplementary Materials

**Supp. Fig. 1.**
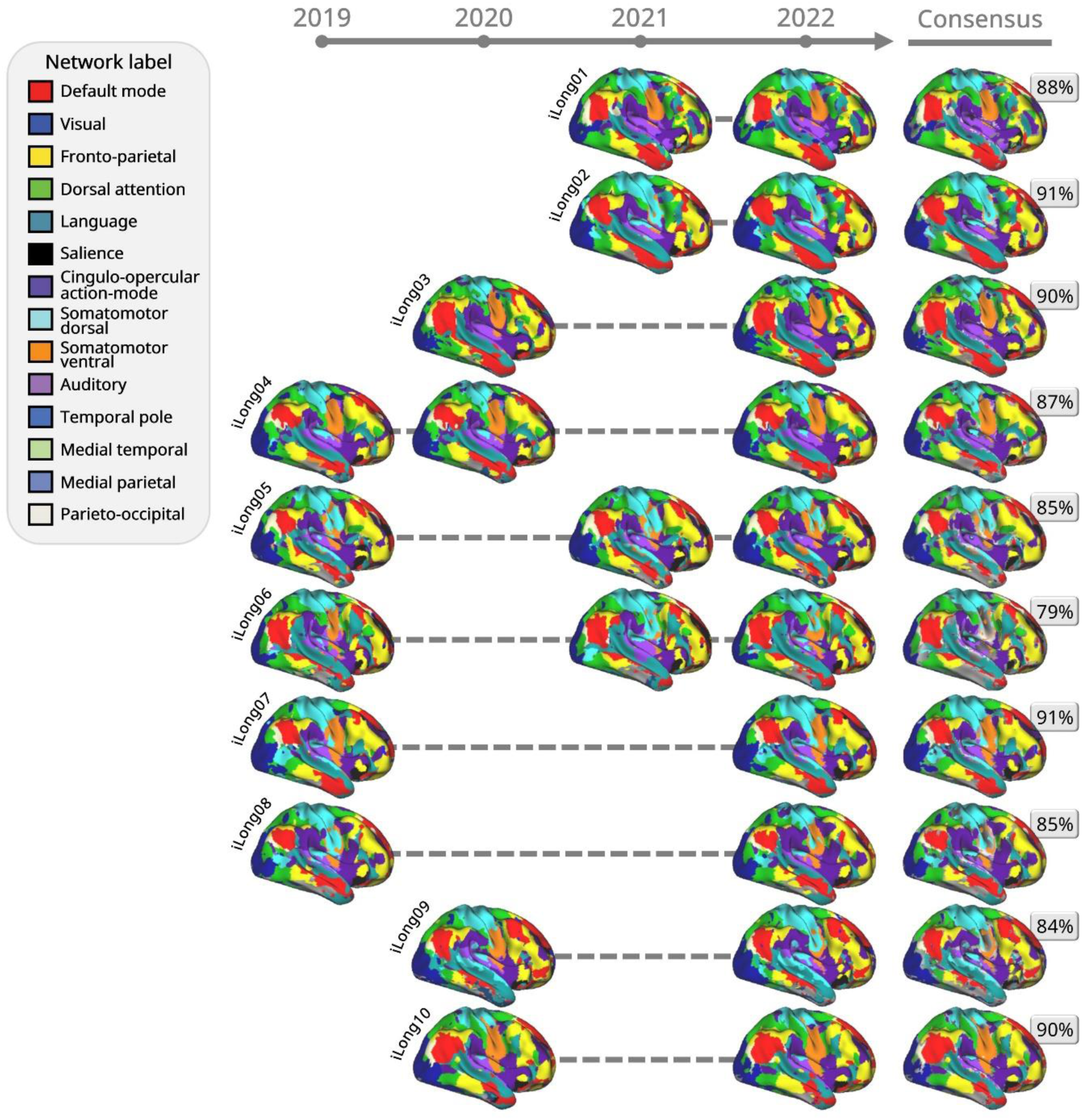
Network consensus and full timeline. For each participant in the Standard Longitudinal Cohort, we mapped functional network organization at the individual level across multiple time points. The resulting consensus maps highlight vertices consistently assigned to the same network across all two or three time points, whereas inconsistent assignments are shown in gray. These maps reveal that network topography is remarkably stable within individuals over time as the percentages of consistent vertices show.

**Supp. Fig. 2.**
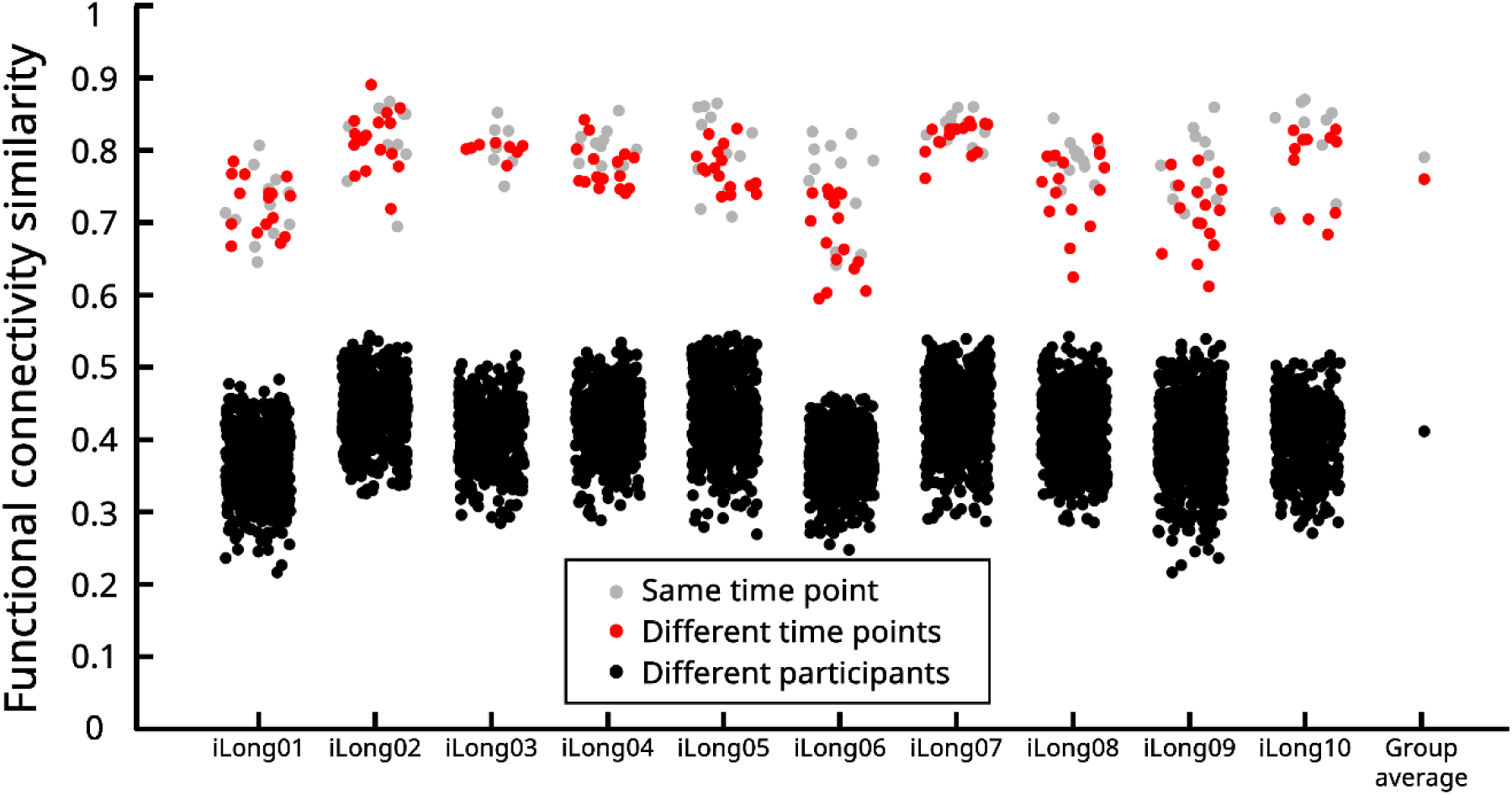
Higher functional connectivity similarity within and across time points than between participants. Volumetric ROI functional connectivity similarity between sessions at the two most distant time points using the Standard Longitudinal data. The gray dots represent similarity within participants within time points (weekly comparisons), the red dots represent comparisons within participants across time points (yearly comparisons), and the black dots represent comparisons between participants. Although weekly comparisons showed the greatest similarity (gray; *M* = 0.79, *SD* = 0.05), yearly comparisons yielded similarly high values (red, *M*= 0.76, *SD* = 0.06) and significantly exceeded between-participant similarity (black dots, *M* = 0.41, *SD* = 0.06; permutation test *p* = 0.002), confirming that functional connectivity patterns are uniquely individual.

**Supp. Fig. 3.**
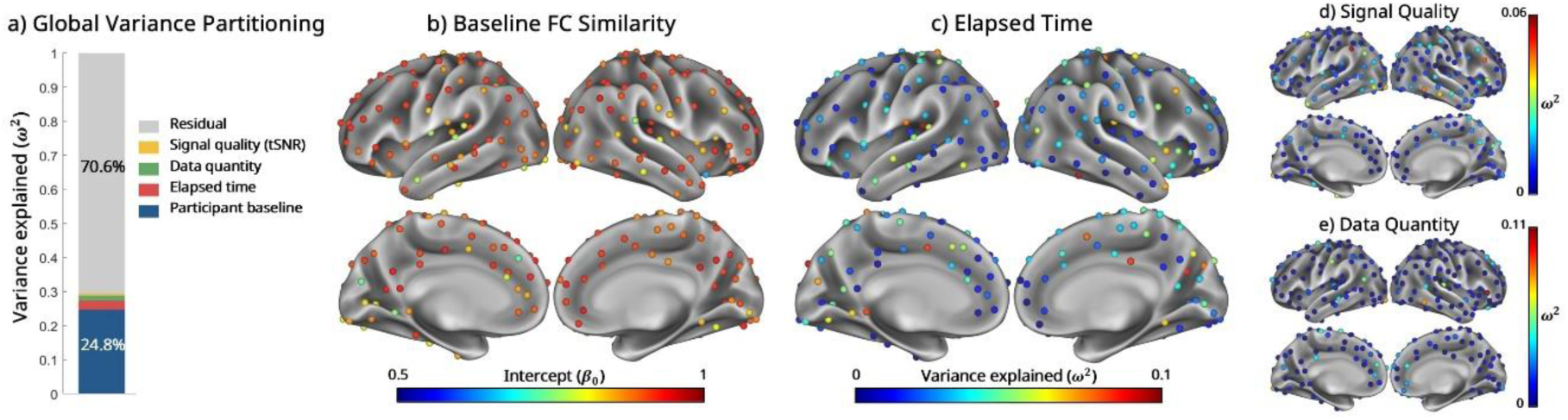
Passage of time explains a small amount of variance in functional connectivity similarity. **a)** Variance explained in a fixed-effects model of between session similarity in functional connectivity, incorporating elapsed time, signal quality, data quantity, and participant baseline. This model accounted for a small proportion (29.4%) of the total variance of functional connectivity (FC) similarity. **b)** The high residual variance (70.6%) in panel a can be explained by high baseline similarity in most regions (Seitzman 300 atlas; shown as foci with subcortical ROIs included in the analyses but omitted here from visualization for clarity). Across most regions, we observed remarkably high similarity (mean intercept *β*_0_= 0.9; the intercept represents the predicted similarity when all predictors are held at their mean as we standardized the values; see *Methods*). **c)** The variance explained (*ω*^2^) by elapsed time was small, supporting our hypothesis that the passage of time had only a subtle effect on FC similarity. The omega-squared values for signal quality **(d)** and data quantity **(e)** were also near zero.

**Supp. Fig. 4.**
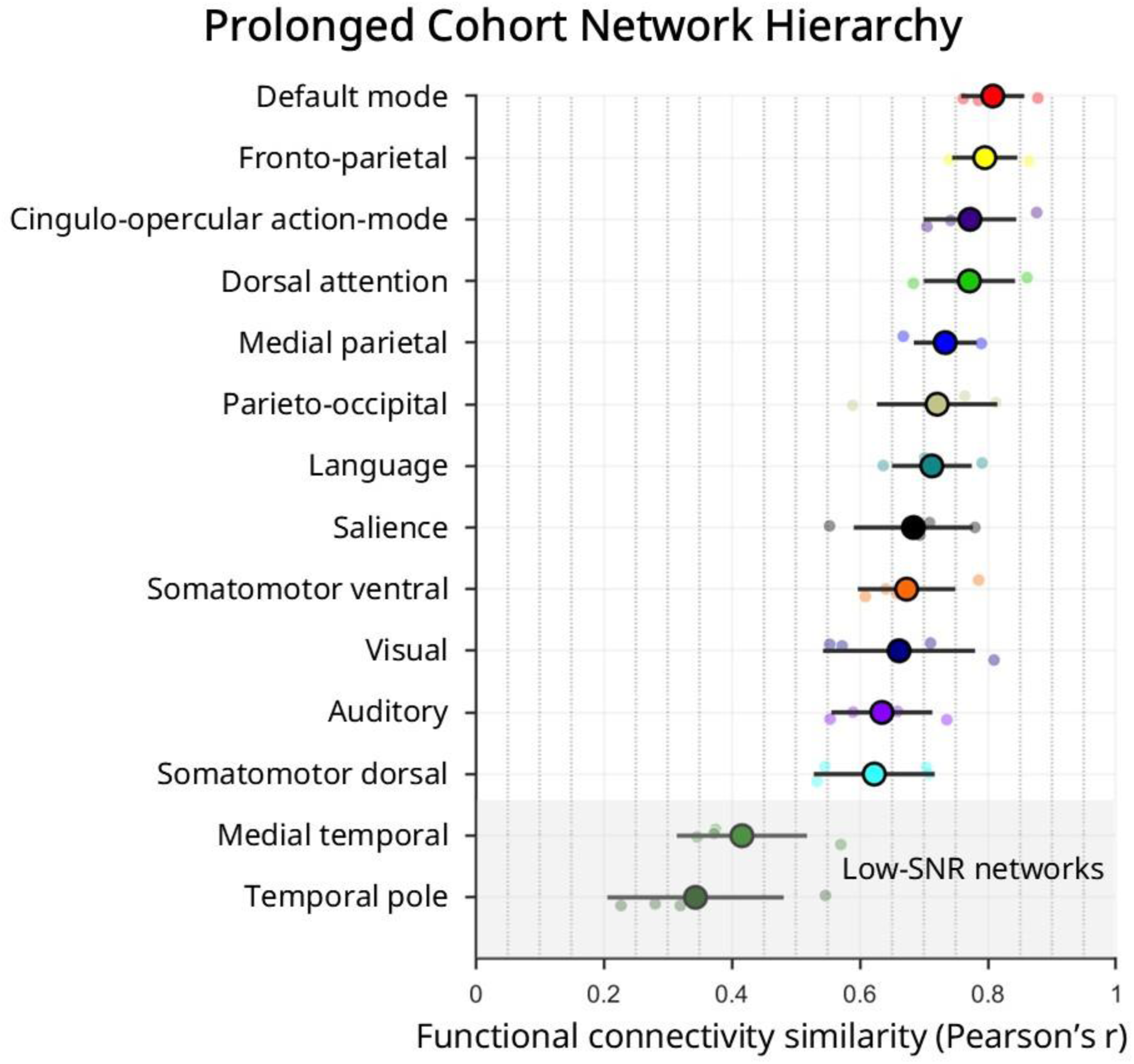
Network stability hierarchy. Network stability rank ordering of the Prolonged Longitudinal Cohort. After generating vertex-wise functional connectivity similarity maps across time points within participants (Fig. 2b), correlation values were averaged within each network (with low-SNR regions censored). Network-level means were then averaged across participants and ranked from highest to lowest stability. The circle indicates the mean similarity within networks across participants with the solid lines showing 95% confidence interval. The ranking is highly similar to that of the Standard Longitudinal Cohort (Fig. 6) where association networks tended to show higher stability than sensorimotor networks (medial temporal lobe and temporal pole are also association networks, but they mostly overlap with low SNR regions).

**Supp. Fig. 5.**
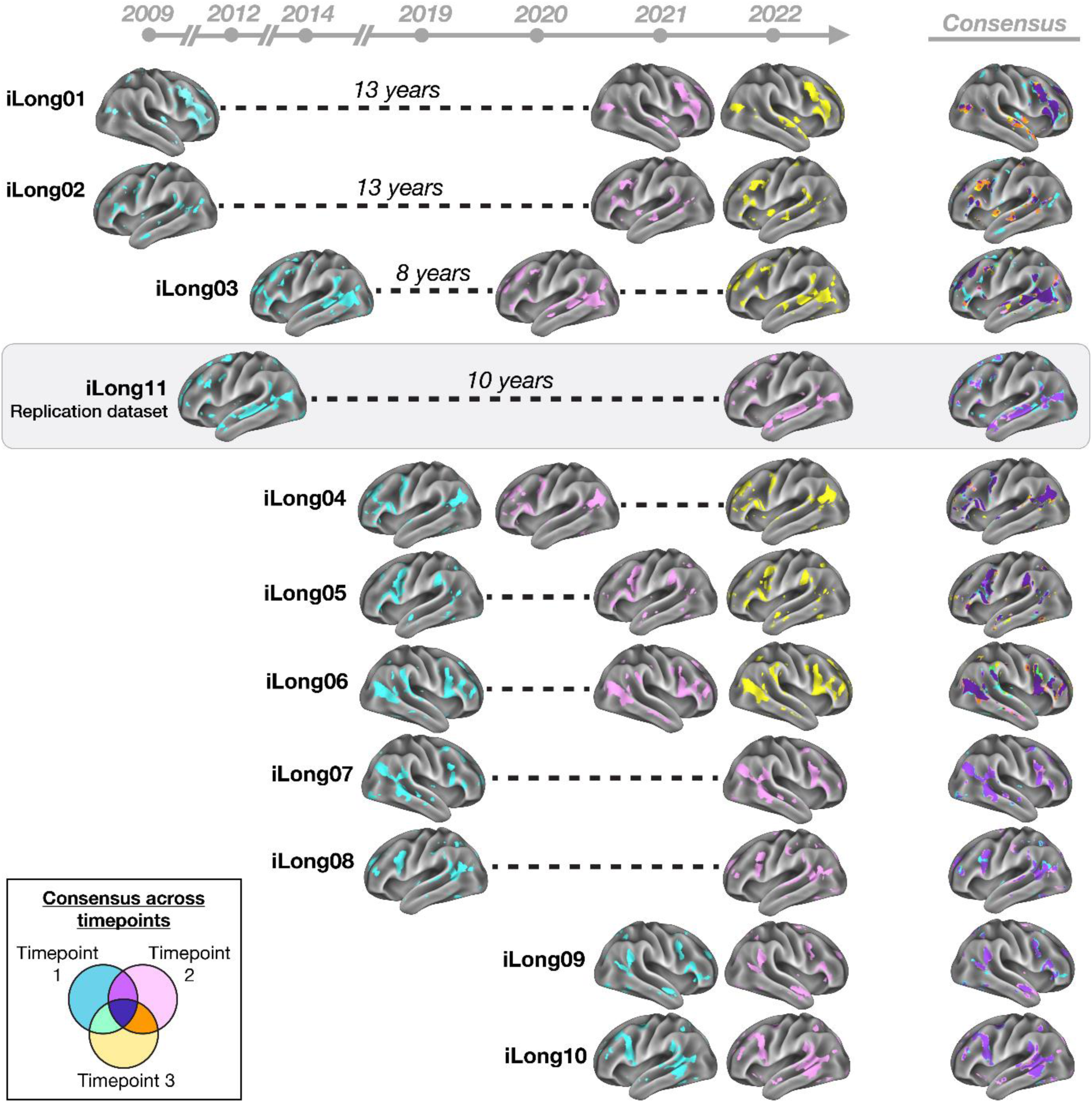
Network variants and full timeline. We defined network variants using bottom 10% individual-to-group functional connectivity similarity threshold at each time point. For each participant, the hemisphere displaying the most variants is shown. The consensus maps display the overlap across time point (color scheme indicated in the Venn diagram). While network variants appeared in different shapes, locations, and sizes across participants, they highly overlapped within participants, even across time spans exceeding a decade.

**Supp. Fig. 6.**
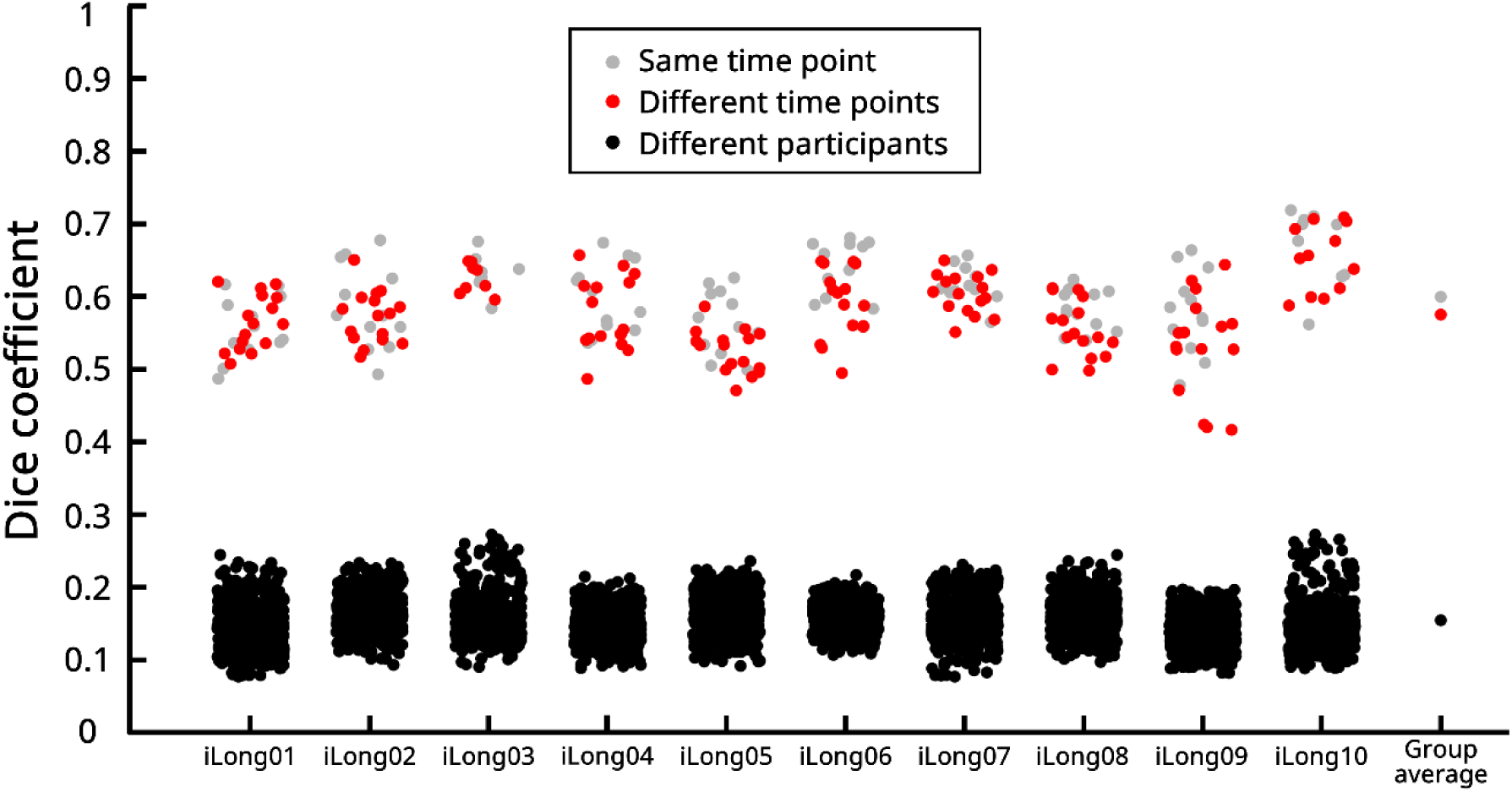
Network variant overlap across weeks and years. Related to Figure 7, this plot illustrates the consistency of network variant locations, comparing sessions within weeks (same time point; gray dots) and across years (different time points; red dots) relative to between-participant similarity (black dots) in the Standard Longitudinal Cohort. Binary maps of network variant locations were compared at two time points using Dice coefficients. Notably, although Dice overlap was highest across weeks (*M* = 0.60, *SD* = 0.05), as expected, it was comparably high across years (*M* = 0.58, *SD* = 0.06), and within-participant yearly comparisons were significantly higher than between-participant comparisons (*M* = 0.16, *SD* = 0.03), *p* = 0.002 (permutation testing). Data amounts were matched, and vertices within low-SNR regions were excluded.

**Supp. Fig. 7.**
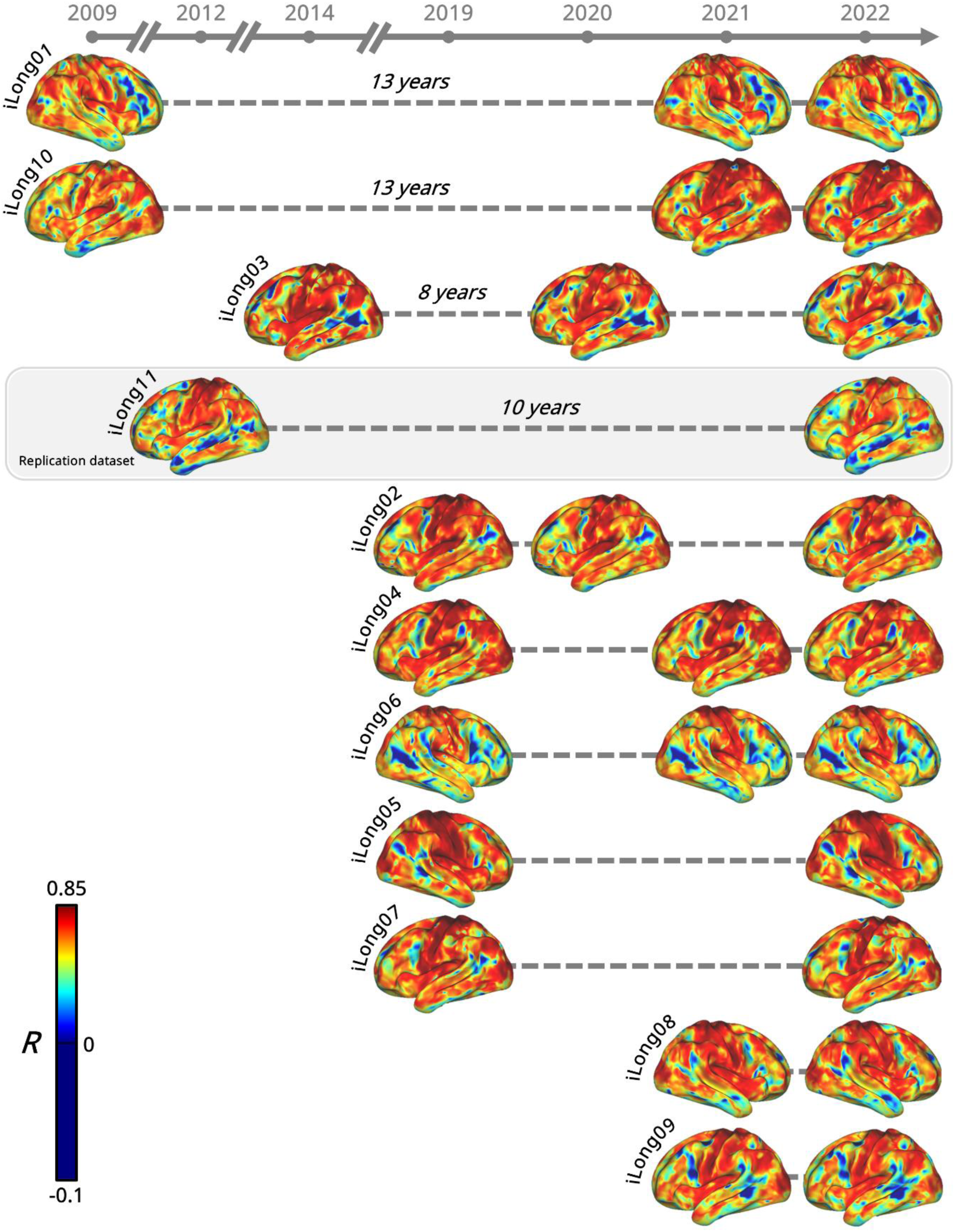
Continuous spatial correlation maps of network variants. We showed in Figure 7 and **Supplementary** Figure 5 network variant locations, using a threshold of below 10%. This shows the full spatial correlation of functional connectivity with an independent group template (WU-120) to capture continuous variation, providing a more nuanced view of consistency across time points. Blue areas, which indicate spatial correlations near zero or negative, appeared in consistent locations across time points.

**Supp. Fig. 8.**
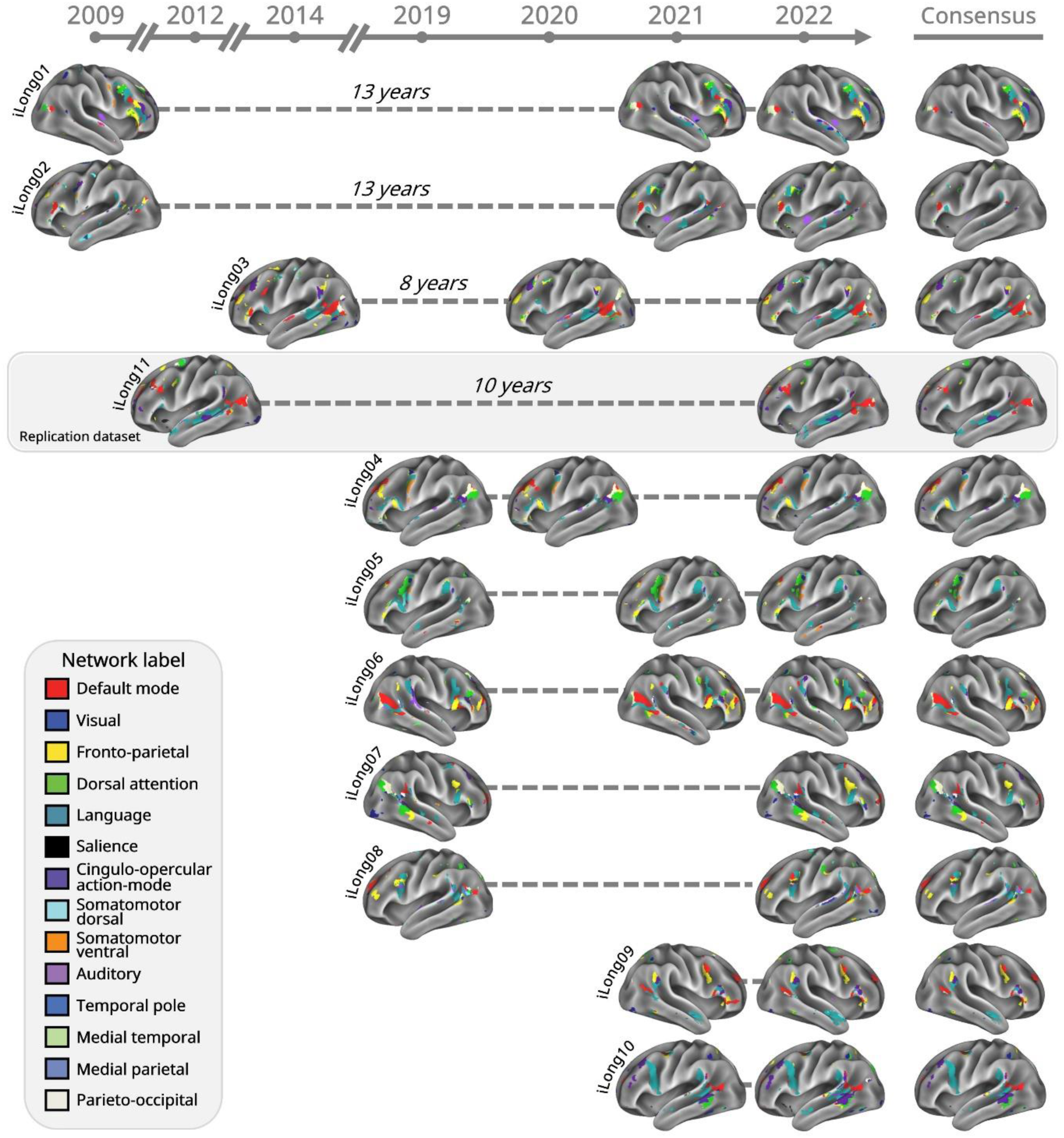
Stable network variant composition. As shown in Figure 7 and **Supplementary** Figure 5, we display here the locations of network variants thresholded at the bottom 10%, but each vertex is also color-coded based on each participant’s template-matching network map (Figs. 2a and 3a). The consensus maps illustrate the network composition of stable variants that appear consistently in the same location across all time points. Notably, the seemingly ectopic variants highlighted in Figures 2a **and 3a** are also evident here.

